# Information theoretic approaches to deciphering the neural code with functional fluorescence imaging

**DOI:** 10.1101/2020.06.03.131870

**Authors:** Jason R. Climer, Daniel A. Dombeck

## Abstract

Information theoretic metrics have proven useful in quantifying the relationship between behaviorally relevant parameters and neuronal activity with relatively few assumptions. However, these metrics are typically applied to action potential recordings and were not designed for the slow timescales and variable amplitudes typical of functional fluorescence recordings (e.g. calcium imaging). The lack of research guidelines on how to apply and interpret these metrics with fluorescence traces means the neuroscience community has yet to realize the power of information theoretic metrics. Here, we used computational methods to create mock action potential traces with known amounts of information. From these, we generated fluorescence traces and examined the ability of different information metrics to recover the known information values. We provide guidelines for how to use information metrics when applying them to functional fluorescence and demonstrate their appropriate application to GCaMP6f population recordings from mouse hippocampal neurons imaged during virtual navigation.

## Introduction

Neurons encode parameters important for animal behavior, at least in part, through the rate of production of action potentials (APs). Evidence for this can be found from electrophysiological AP recordings of orientation tuning in the visual system (Hubel and Wiesel, 2009), chemical sensing in the olfactory system (Leveteau and MacLeod, 1966; Wachowiak and Shipley, 2006), and spatial encoding in the hippocampus (O’Keefe, 1976). Key to deciphering the neural code, therefore, is defining metric to quantify the relationship between behavioral parameter spaces and a neuron’s spiking rate. There are many metrics used for quantification, and are often used to compare neural responses across conditions or in neurons with complex responses. The underlying assumptions of the different metrics then become important factors to consider when determining which one to use.

Information theory is growing in popularity in the neuroscience community, largely because it provides a means to quantify rate coding with relatively few assumptions. One useful information theoretic measure is mutual information (MI), which is typically measured in bits per unit time, and describes the increase in predictability of the neural response when behavioral parameters are known. Formally, mutual information is the information about one variable that can be extracted from another, such as the information about behavior that can be derived from observing neural activity. Mutual information can be applied to neurons with widely varying response properties because it:

1. Is a nonlinear metric, not requiring the linearity assumptions of correlation metrics (e.g. (Grubb and Thompson, 2006; Hinman et al., 2016; Jiaying Tang, 2015; Kropff et al., 2015),
2. Does not assume a response shape, as is typical with Gaussian field mapping metrics (e.g. Kraus et al., 2015; Soo et al., 2011) or metrics using exponential or polynomial curve fitting (Hinman et al., 2016; Jiaying Tang, 2015), and
3. Uses the full time trace or shape of the mean response profile, rather than defining receptive fields with thresholding (e.g. Harvey et al., 2009; Niell and Stryker, 2008; Pastalkova et al., 2008).

However, MI can be nontrivial to estimate from neural and behavioral recordings and its estimation is an ongoing area of research (Belghazi et al., 2018; Gao et al., 2017; Kraskov et al., 2004; Timme and Lapish, 2018).

Here we focus on the most widely used estimator of MI in neuroscience, the SMGM estimator developed by Skaggs, McNaughton, Gothard and Markus (Skaggs et al., 1993), though as a point of comparison, we also consider the Binned Estimator (Timme and Lapish, 2018) and a separate technique developed by Kraskov, Stogbauer and Grassberger (KSG, Kraskov et al., 2004). The Binned Estimator estimates the joint probability distribution using a 2D histogram of neural response vs. behavioral variable; this transforms continuous variables into discrete values (Timme and Lapish, 2018). KSG estimates mutual information by examining the distance between data-points in the neural activity-behavioral parameter space. The SMGM estimator, on the other hand, relies on the assumption that AP firing follows an inhomogeneous Poisson process. The SMGM estimator therefore requires binning of only the behavioral variable(s), in contrast to the Binned Estimator. The profile of firing rates vs. behavioral variable is then used to estimate the MI.

The relative simplicity of the SMGM estimator has added to its popularity and widespread use in neuroscience applications for estimating behavioral information contained in single unit AP recordings. This metric has proven useful in quantifying rate coding in place cells (Knierim et al., 1995; Lee et al., 2006; Markus et al., 1995; Poucet and Sargolini, 2013), complex spatial responses of hippocampal interneurons (Frank et al., 2001; Wilent and Nitz, 2007), odor sequence cells (Allen et al., 2016), time cells (MacDonald et al., 2013), head direction cells (Stackman and Taube, 1998), speed cells (Fyhn et al., 2002), and face differential neurons (Nguyen et al., 2014, 2013), and has been used across multiple different species (Hazama and Tamura, 2019; Mankin et al., 2019; Yartsev and Ulanovsky, 2013). Furthermore, as a single neuron metric it provides statistical power for comparisons. Thus, it has been used to quantify differences in rate coding across different brain regions (Simonnet and Brecht, 2019) and across experimental interventions such as lesions (Calton et al., 2003; Liu et al., 2004), inactivations (Brandon et al., 2011; Hok et al., 2013; Huang et al., 2009; Koenig et al., 2011), and applications of drugs (Newman et al., 2014; Robbe and Buzsáki, 2009). Further, it has been used to examine differences in encoding across different behaviors (Aronov and Tank, 2014; Park et al., 2011; Zinyuk, 2000), and disease states (Fu et al., 2017; Gerrard et al., 2008; Zhou et al., 2007). SMGM information is often normalized from measuring bits per unit time to instead measure bits per AP. This creates a measure sensitive only to the selectivity of a neuron, and not its average firing rate. Thus, SMGM is a powerful tool for measuring the neural code in electrophysiological recordings of APs.

The power of MI estimators has yet to be fully exploited by the neuroscience community. For example, the estimators have not yet been widely used to compare encoding properties of large numbers of genetically identified neurons, or to quantify information content of other discrete signaling events such as synaptic inputs; both of which are difficult to study using electrophysiological methods. *In vivo* imaging of functional indicators has emerged as an important tool, largely because it possesses these capabilities. For example, using fluorescent calcium indicators, the functional properties of large populations of neurons can be simultaneously recorded in rodents (Dombeck et al., 2007; Radvansky and Dombeck, 2018; Sheffield et al., 2017; Stirman et al., 2016; Stringer et al., 2019; Ziv et al., 2013) zebrafish (Ahrens et al., 2013), or invertebrates such as *C. elegans* (Nguyen et al., 2016) and *Drosophila* (Keller and Ahrens, 2015; Mann et al., 2017). Furthermore, *in vivo* imaging can assure the genetic identity of the recorded neurons (Jing et al., 2018a, 2018b, 2018c; Khoshkhoo et al., 2017; Sheffield et al., 2017) and can access subcellular structures, allowing for functional recordings from synapses and dendrites using different functional fluorescent indicators (e.g. Jing et al., 2018d; Marvin et al., 2019, 2018; Scholl et al., 2017; Sheffield et al., 2017; Sheffield and Dombeck, 2015).

However, these indicators generate signals that are different from the underlying quantal events. For example, somatic calcium indicators reveal intensity variations that are correlated with somatic AP firing rates but are a smoothed and varying amplitude version of the AP train. This transformation from AP train to fluorescence trace is an active area of research (Dana et al., 2018; Éltes et al., 2019; Greenberg et al., 2018), but it is often approximated by convolving the AP train with a kernel, which defines the indicator’s response to a single AP. The shape of the kernel is a function of the indicator expression level, intracellular calcium buffering, amount of calcium influx, efflux rates, background fluorescence, resting calcium concentration, and other factors. When measured in pyramidal neurons, average kernels typically take the shape of a sharp increase in fluorescence followed by an exponential decay to baseline (Chen et al., 2013; Dana et al., 2018; Pachitariu et al., 2018; Park et al., 2013; Yaksi and Friedrich, 2006). Therefore, while functional fluorescence imaging and information theoretic quantification may prove to be a powerful new combination of tools to study neural correlates of behavior, it is critical to remember that functional fluorescence signals represent altered versions of the underlying physiological events.

Caution is then needed when applying information metrics to continuous functional fluorescence traces, yet the imaging community is already beginning to use information metrics, particularly SMGM. This metric has been applied to somatic calcium responses to compare the information content of the same neurons across different behavioral epochs (Heys and Dombeck, 2018), across different populations of neurons in different brain regions (Hainmueller and Bartos, 2018), across different genetically identified neural populations (Khoshkhoo et al., 2017), or to examine encoding by subcellular structures (Rashid et al., 2020), or to classify the significance of encoding particular parameters by individual neurons (Kinsky et al., 2018; Mau et al., 2018; Rashid et al., 2020).

However, it is essential to recognize some of the assumptions underlying these information metrics are violated by functional florescence recordings. All three metrics (SMGM, KSG and Binned Estimation) assume stationarity in the neural response, which is violated by the elongated time responses and relatively slow fluctuations of the fluorescence intensity of the reporters. When applied to spiking data, there is also a change in units: rather than AP counts, functional fluorescence traces are typically plotted in units of florescence change with respect to baseline (ΔF/F). One possible solution to these issues would be to deconvolve calcium traces to recover APs; however, deconvolution is an active area of research, and the accuracy of these methods has recently been questioned (Evans et al., 2019). Ideally, the calcium traces could be used directly to measure spiking information, without the need for such an in between, potentially error inducing, step.

Quantifying the effects of the above violations on measurements of information using functional fluorescence recordings with an analytical solution is particularly challenging with behaviorally modulated neural recording data. However, a more tractable means of quantifying the effects would be to use a simulation study to measure the induced biases and changes in measurement quality (Morris et al., 2019). This strategy makes use of pseudo-randomly generated AP traces and has the advantage that the ground truth parameters of the simulations are known, while variability due to behavior and other features can be incorporated (Climer et al., 2015, 2013; Cohen and Kohn, 2011; Østergaard et al., 2018).

To provide the field with guidelines for the use of information metrics applied to functional fluorescence recording data, we used computational simulation methods to create a library of ten thousand mock neurons whose spiking output carry an exact, known (ground-truth) amount of information about the animal’s spatial location in its environment. We used real behavioral data (available at https://doi.org/10.7910/DVN/SCQYKR) of spatial position over time from mice navigating in virtual linear tracks and then simulated the spatial firing patterns of the mock neurons using an inhomogeneous Poisson process framework (Brown et al., 2003; Climer et al., 2013; Paninski, 2004). We then simulated fluorescent calcium responses for each neuron in each session by convolving the AP trains with calcium kernels for different indicators, primarily GCamp6f (Chen et al., 2013), and then we added noise. MI metrics (between spatial location and the neural signals) were then applied to the spiking or fluorescence traces to quantify the performance of the metrics for estimating information. We provide a user toolbox (found at https://github.com/DombeckLab/infoTheory), which consists of Matlab functions to generate libraries of model neurons with known amounts of information, to generate spiking or fluorescence time-series from those model neurons, and to estimate neuron information from real or model spiking or fluorescence time-series datasets using the three metrics considered here (SMGM, Binned Estimator, KSG). We focused on testing the performance of the SMGM method, and then compared its performance to the Binned Estimation and KSG methods, which do not have the underlying Poisson assumption required for the SMGM approach. We also applied a deconvolution algorithm to test its performance. We then applied this analysis to real datasets of hippocampal neuron populations from mice navigating in virtual linear tracks. We quantified the spatial information content of the populations and then performed Bayesian decoding of mouse position from different information containing subsets of this population. Interestingly, we found that the population quantile with the lowest information values were still able to decode mouse position to the closest quarter of the track. Thus, we provide new findings about the neural code for space that were made possible by the information metrics and guidelines that we introduce here.

The SMGM method applied directly to the mean ΔF/F intensity map appeared to best recover the ground truth information. We provide guidelines for the use of the SMGM metric when applied to functional fluorescence recordings and demonstrate the appropriate application of these guidelines to GCaMP6f population recordings from hippocampal neurons in mice navigating virtual linear tracks.

## Results

### The SMGM information metrics

Here we review the derivation of the SMGM information metrics and the underlying assumptions. For illustrative purposes throughout this manuscript, we use the example of spatial encoding in which the firing pattern of neurons carry information about the animal’s location along a linear track; however, the derivations, equations and conclusions generalize to encoded variables over other domains and dimensionalities.

Consider a random variable X representing the positions an animal might take, with *x* being its value measured at one time sample. The positions are subdivided into N spatial bins, such that *x* can take on the values {1,2,…, *N*}. For our analyses, *N*=60. Consider a random variable Y representing the number of APs a neuron might fire, where *y* is the count measured within a time sample. *y* can take on the values of {0,1,…, +∞}. X and Y are both discrete. If X and Y both obey the assumption that each time sample is independent (i.e. they are stationary), then the mutual information (I, in bits per sample) between X and Y is expressed as follows:

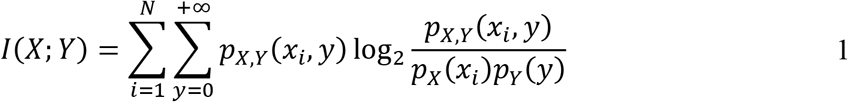

Where *p*_x_(*x_i_*) is the (marginal) probability that the animal is in the *i*^*th*^ spatial bin during a time sample, *p*_Y_(*y*) is the probability that the neuron fires *y* APs in the time sample, and *p*_X_,_Y_(*x_i_*, *y*) is the joint probability that the neuron fires *y* APs and is in the *i*^*th*^ bin. Recall that *p*_X_,_Y_(*x_i_*, *y*) = *p*_Y|X_(*y*|*x_i_*)*p*_X_(*x_i_*), where *p*_Y|X_(*y*|*x_i_*) is the conditional probability that the neuron fires *y* APs given that the animal is in the *i*^*th*^ spatial bin. We can thus rewrite Equation 1 as follows:

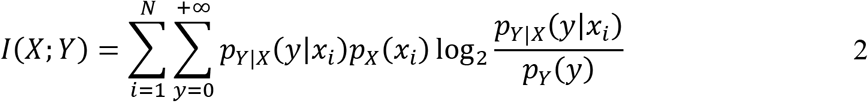

With the further assumption that the firing of the neuron follows Poisson statistics, we can then estimate the mutual information as follows: let the AP rate (AP/s or Hz) in a single bin be λ_*i*_, and the average across the session be 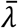. For an arbitrarily small time window Δ*t*, the probability that an AP occurs in that window is Pr(*Y* = 1|*x* = *i*) = λ_*i*_ Δ*t*, with the probability that an AP occurs regardless of position as 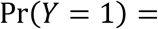 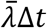. We can thus rewrite Equation 2 as:

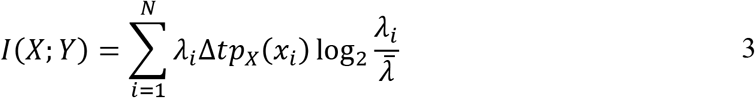

By integrating over one second 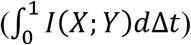 we obtain the first key SMGM metric for spatial information as measured by AP firing, which is in units of bits per second:

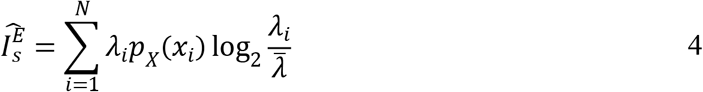

For notation, we will use a carrot (^) to indicate an information value that is measured from experiment, the superscript (E in this case) to show the source of the data, and a subscript to show the units/formula used (bits per second in this case). Thus, 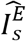 is the information measured via electrophysiology in bits per second. This metric is linearly dependent on the average firing rate of the neuron, and this dependence is often removed through normalization by the average firing rate to obtain the second key metric of spatial information as measured by AP firing, which is in units of bits/(second*Hz), or more commonly, bits per AP:

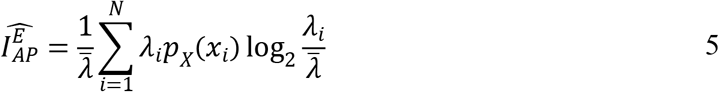

Therefore, these two key metrics of spatial information are defined completely by quantities that can be experimentally measured: the mean firing rate 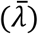 from the AP counts over the duration of the recording, the AP firing rate in the *i*^*th*^ bin from the average rate map (λ_*i*_*i*), and the probability that the animal is in the *i*^*th*^ spatial bin from the normalized occupancy map (*p*_X_(*x_i_*)). The quantity of and noise in these measurements affects the quality of the metric: in particular, undersampling due to low firing rates or low trial counts induces a substantial positive bias (Treves and Panzeri, 1995).

In the derivation of these metrics, there are two key assumptions that are violated by functional fluorescence recordings. First, the recordings do not follow Poisson statistics: instead of discrete counts of APs (*y*), the functional fluorescence traces consists of a continuous relative change in fluorescence (Δ*F*/*F*), and instead of a firing rate map (λ_*i*_*i*) measured in Hz, average intensity maps in units of Δ*F*/*F* are generated. The stationarity assumption is also violated: due to the slow decay, a time sample of the fluorescence traces depend on the previous samples. The violation of these assumptions by functional fluorescence recording will affect the precision and induce biases in the SMGM information metrics. Since these effects have not previously been addressed or quantified, we measured these biases here using a simulation study.

### Building a ground truth library of 10,000 neurons with known values of information

To create a neuron with a known, ground truth information value, it was necessary to generate a continuous (i.e. infinitesimally small bins) rate map (λ(*x*)) matching the desired information. To do this, we first normalized the track length to 1 and assumed the animal’s occupancy map to be spatially uniform 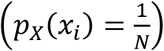. We then created an exponentiated cubic spline with 5 randomly positioned nodes (Figure 1A) to build a starting continuous map of the normalized instantaneous firing rate, 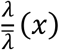, with the integral normalized to 1. We calculated the ground truth amount of information in bits per AP as follows:

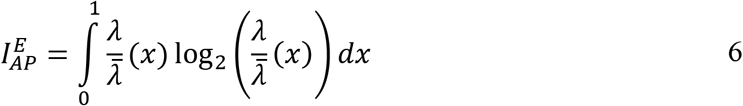

The locations of the 5 nodes were then systematically varied [see Methods] to minimize the squared error between the value calculated in Equation 6 and a target amount of information (Figure 1A,B), in the end resulting in a mean error of 5.1*10^-9 bits/AP and a mean absolute error of 1.5*10^-7 bits/AP. The rate map at this convergence point was used for further analysis. This procedure was repeated to generate 10,000 mock neurons with a range of (known and ground truth) information values. Note that the value in Equation 5 cannot be higher than when all the APs arrive in one spatial bin; the rate in that bin is 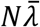. If we assume uniform occupancy 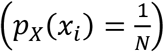, then the maximum measureable information is log_2_ *N*, in our case, 5.9 bits/AP with *N*=60 bins. Thus, the information values considered here range between 0 and 6 bits/AP (Figure 1C). We chose a mean firing rate 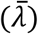 for the neurons between 0.1 and 30 Hz, a range observed for a variety of different cortical and hippocampal neurons during behavior (Buzsáki and Mizuseki, 2014; DeWeese et al., 2008; O’Connor et al., 2010; Roxin et al., 2011; Shafi et al., 2007). From Equations 4–5, the ground truth information in bits per second is 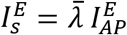. 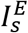 for these choices resulted in ground truth information values between 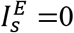 and 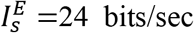 (Figure 1C). Example low 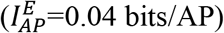 and mid 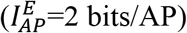 rate maps are shown in Figure 1D.

**Figure 1.**
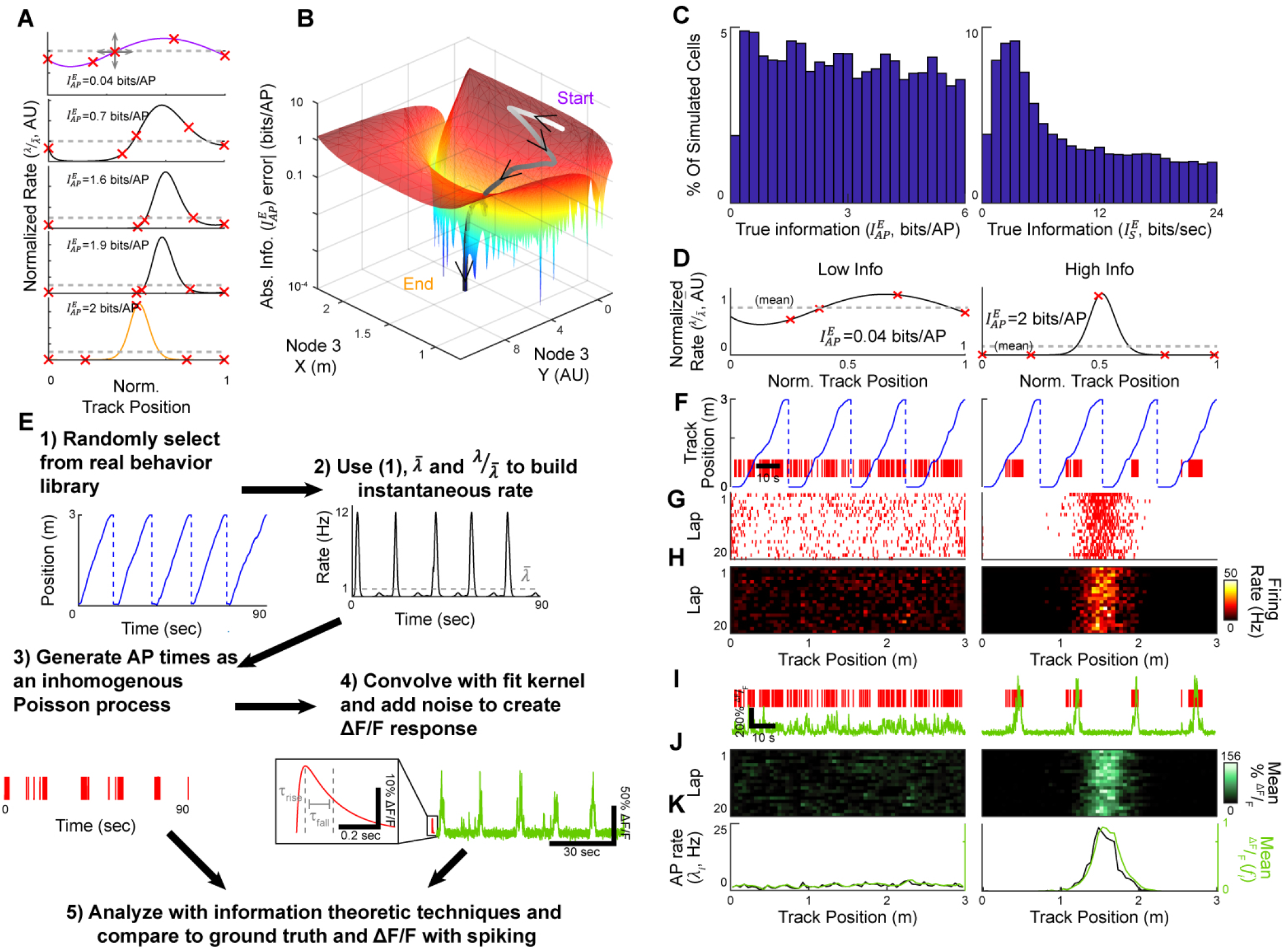
Procedures for generating a library of 10,000 neurons with known amounts of information. (A) Five splines with a gradient of ground truth information 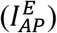 representing the steps in generating a continuous rate map (λ_*i*_(*x*)) ma tching th e de sired ta rget information, in this case 2 bits/AP. Re d X’ s indicate control nodes that were moved to change the shape of the spline and minimize the squared error to the target information. (B) Cross section of the error surface around the solution point as a function of the position of node 3, and the trajectory taken by the solver to minimize the error and arrive at the target. (C) Histograms of ground truth information resulting from repeating the procedure in A-B 10,000 times to target a range of ground truth information values in bits per second 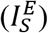 and bits per AP 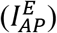. (D) Splines representing λ_*i*_ (*x*) at the solution point for a low (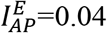 bits per AP, left) and high (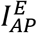, 2 bits per AP, right) information neuron. (E) Steps to generate mock AP and functional fluorescence data. (1) An example real behavior trace from a mouse running on a linear track that was used to generate the simulated spiking. (2) The behavior in combination with the rate maps generated in A-D were used to generate an instantaneous firing rate trace. (3) The instantaneous rate was used to pseudorandomly generate APs, as shown in this mock raster. (4) The AP raster was convolved with the GCaMP6f kernel (red, inset) and noise was added to generate a mock 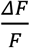 trace. (5) Large numbers of these traces were generated and used to assess the effects of many simulation parameters on the estimators. (F-L) Spiking and fluorescence activity patterns generated from the example simulated neurons shown in D and using a mean firing rate of 1 Hz. (F) Behavioral trace in blue with AP raster shown in red. (G) Lap-by-lap raster of the neurons’ firing vs mouse track position. (H) Lap by lap binned, firing rates vs mouse track position for the neurons (I) AP raster (red) and mock calcium traces for the same behavioral period shown in F. (J) Lap by lap mean binned fluorescence vs mouse position for the neurons. (K) Binned average firing rate (λ_*i*_*i*, black) and fluorescence intensity (*f_i_*, green) maps for the two neurons. These maps were used for information analyses.

These rate maps provided a basis for generating mock AP firing data (and functional fluorescence data, see below). Under real experimental conditions, recording duration and bin sizes are finite and animal occupancy maps (*p*_X_(*x_i_*)) are not spatially uniform. These experimental limitations add error to the estimate of a neuron’s ground truth information value. Therefore, in order to accurately re-create these limitations in our simulation study, we used real behavior datasets from head-restrained mice running along a 3 m virtual linear track for water rewards (acquired as in Sheffield et al., 2017; Sheffield and Dombeck, 2015). Unless otherwise indicated, all values reported will be the mean±standard deviation. We selected at random from a library of 574 behavior sessions from mice navigating along familiar tracks and concatenated and truncated these sessions to create behavior sessions uniformly sampled up to 60 minutes in duration (average 30.2±17.1 minutes), resulting in an average 132 ± 71.2 laps per session and an average running speed of 19.3 + 3.87 cm/sec (Figure 1E.1). This behavior, the average firing rate 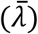, and the normalized rate map 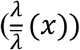 from the mock neurons were used to create an instantaneous firing rate trace (Figure 1E.2), sampled at 1 kHz, from which AP times were generated assuming Poisson firing statistics (Figure 1E.3). An example mock of spiking in response to behavior for low (0.04 bits/ AP) and mid (2 bits/AP) information neurons can be seen in Figure 1F-H. From these spiking responses, we then generated mock fluorescence traces by convolving the raster with a double-exponential kernel matching the rise and fall times for GCaMP6f (Chen et al., 2013, Figure 1E.4) and adding random Gaussian noise to model shot noise. Mock fluorescence traces for the two example neurons in Figure 1F-H can be seen in Figure 1I-J. The mock AP and fluorescence traces were used to create session mean spatial maps – of binned firing rate (λ_*i*_ in Hz) and change in fluorescence (in Δ*F*/*F*), for information analyses (Figure 1K). By repeating this process, we built a large dataset of spiking and fluorescence traces, generated from our library of mock neurons with known amounts of information and using real animal spatial behavior. With tens of thousands of these mock neuron recordings, we could then assess the effects of many simulation parameters on the information values determined from the metrics including firing rate, session duration, fluorescence kernel shape, and ground truth information value.

### Quantification of the accuracy and precision of the SMGM bits per second metric using functional fluorescence recordings

We first applied the SMGM bits per second metric 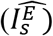 to our mock AP recording traces to verify that they can recover our ground-truth information values given finite recording durations and bin sizes, and non-uniform animal occupancy maps (*p*_X_(*x*)). Figure 2A shows three mock neurons with ground truth information values of 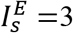, 15 and 23 bits/sec. When the SMGM bits per second metric 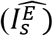 was applied to the AP traces from these example neurons, the information was well recovered, with 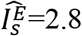, 15 and 24 bits/sec, respectively. The results from these examples also held across the full 10,000 mock neuron library (Figures 2B-D), as a linear fit (y-intercept = 0.093±0.040, intercept p=4.6*10^−6^bits per second and slope = 0.97 ±0.0030, slope p<<0.01)explained nearly all the variance (R^2^= 0.97), the average error was 0.22±1.25 bits per second (1.0±0.69% error) and the absolute error was 0.64±1.05 bits per second (8.4±0.69% error). There is a substantial positive bias for the lowest firing rates and smallest number of trials (Figure 2-figure supplement 1A-B) which has been previously well characterized (Treves and Panzeri, 1995), with average errors exceeding +10% for less than 6 minutes of recording, mean rate under 0.6 Hz, and under 11 trials. Thus, the SMGM bits-per-second metric 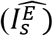 recovers the ground-truth information well using AP recordings, with the only error coming from finite recording time and variable animal behavior.

**Figure 2.**
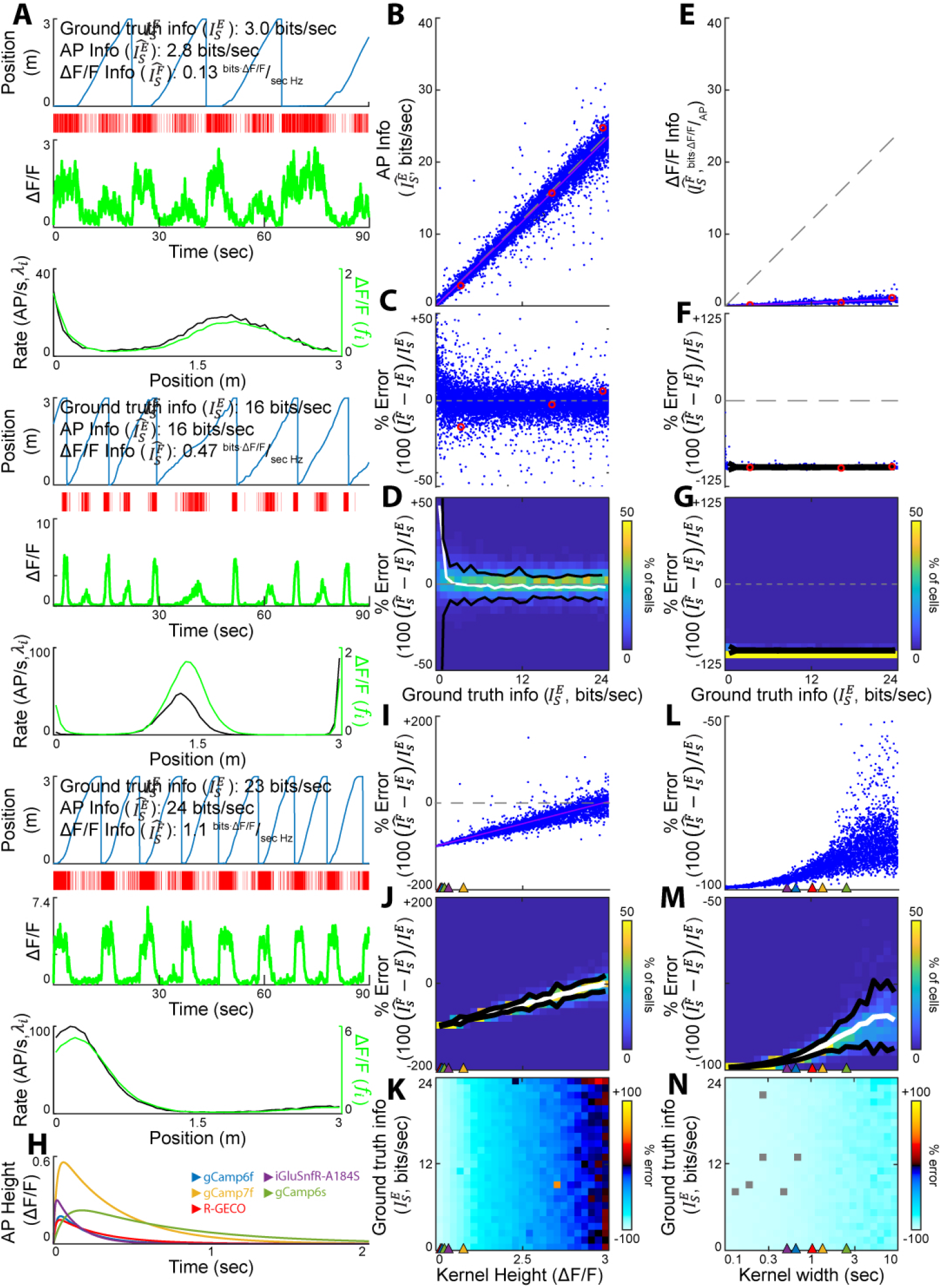
Quantification of the precision of the SMGM bits per second metric using APs or functional fluorescence recordings. (A) Three representative mock neurons spanning the range of ground truth information values in bits per second 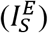. From top to bottom for each: mouse track position vs time, AP raster, fluorescence calcium trace (green), and firing rate map (λ_*i*_, black) and change in fluorescence map (fi, green). (B-D) The ground truth bits per second values are well recovered when measured from AP traces. (B) Information measured from AP data using the SMGM bits per second metric 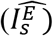 vs ground truth information 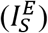. Each dot is a single mock neuron, the gray dashed line is the unity line (perfect measurement), the pink line is the line of best fit. Red circles show the examples in A. (C) Percentage error for the information measurements shown in B. (D) Heat map of percentage error measurements shown in C. Black lines are 2 standard deviations, the white line is the mean. (E-G) Effects of applying the SMGM bits per second metric to fluorescence traces. (E) Information measured from mock GCaMP6f traces using the SMGM bits per second metric 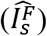 vs ground truth information 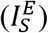. (F) Percentage error for the information measurements shown in E. (G) Heat map of percentage error measurements shown in F. (H) Representative mock kernels mimicking responses from different indicators. (I-K) The effect of kernel height on estimating ground truth information 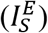 using the SMGM bits per second metric 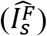. Kernel height for the kernels shown in H are indicated by colored triangles (I) Percentage error as a function of kernel height (J) Heat map of percentage error measurements shown in I with mean (white) and 2 standard deviations (black). (K) The average percentage error as a function of kernel height and ground truth information in SMGM bits per second 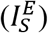. (L-N) The effect of kernel width on estimating ground truth information 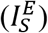 using the SMGM bits per second metric 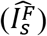. Kernel widths for the kernels shown in H are indicated by colored triangles. (L) Percentage error as a function of kernel width (M) Heat map of percentage error measurements shown in L with mean (white) and 2 standard deviations (black).(N) The average percentage error as a function of kernel width.

We next discuss the changes to the SMGM bits per second metric 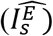 commonly used for application to functional fluorescence traces (Hainmueller and Bartos, 2018; Heys and Dombeck, 2018), and explore the implications of these changes. Most simply, the mean firing rate 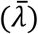 and the mean firing rate in a spatial bin (λ_*i*_) are replaced by the mean change in fluorescence 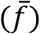 and the mean change in fluorescence in a bin (). Making these substitutions in Equation 4 results in the information as measured by functional fluorescence:

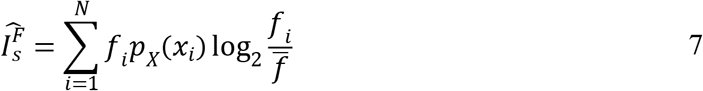

The fluorescence map *f_i_* differs from the firing rate map λ_*i*_ in two ways. First, the fluorescence map is approximated by the firing rate map scaled by a factor *c*, dependent on the height and width of the kernel and measured in units of 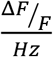, and second it is smoothed by the kernel (Figure 1E4). If we discount the latter for a moment and focus on the scaling, *f* ≈ *c*λ, we can see that substituting λ with *c*λ in Equation 4 results in 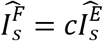. The units for 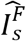 are no longer in bits per second, as it has previously been reported (Hainmueller and Bartos, 2018), but are instead in units of 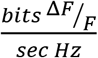 or 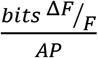 which are difficult to interpret (see *Guidelines for application of information theoretic metrics to functional fluorescence imaging data* section for further discussion). The effect of smoothing is difficult to analytically quantify since it both alters *c*by changing the average intensity and distorts the firing rate map. Therefore, to fully quantify the impact of convolving an AP recording with a functional fluorescence kernel on recovering ground truth information, we used our mock fluorescence traces.

We applied a GCaMP6f modeled kernel to the 10,000 mock AP traces to generate 10,000 mock fluorescence calcium traces. Figure 2A shows the fluorescence traces generated from three mock neurons with ground truth information values of 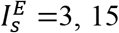, and 23 bits/sec. The effects of the convolution can be seen in the differences in scaling and shape between the fluorescence maps *f_i_* and the firing rate maps λ_*i*_. When the fluorescence metric 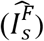 was applied to the fluorescence traces from these example neurons, the information recovered was 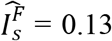, 0.47 and 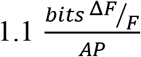, respectively, indicating deviation from the ground truth information values assuming the units are comparable. The results from these examples also held across the full 10,000 mock neuron library (Figures 2E-G), as there was a clear scaling of the ground truth information and a consistent overestimation with a mean error of −11.1±6.7 AU (−96.0±1.3% error). The best-fit line of the measured information 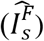
versus the ground truth information 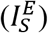 had an intercept near 0 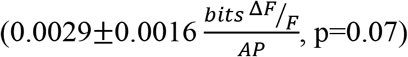. The slope of this fit was 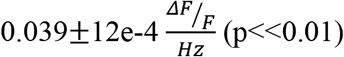, which provides a measure of the scaling factor (*c*). This error was not corrected for with denser sampling: it remained consistent even at high firing rates and many trials (Figure 2-figure supplement 1C-D). In addition to this scaling effect caused by *c*, smoothing of the rate map could induce nonlinearity in the relationship between 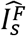 and 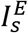. To test for such an effect, we fit the measured information in Figure 2E with a saturating exponential and compared the fits using a likelihood ratio test: the exponential did not significantly improve the fit (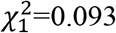, p=0.76), which indicates that smoothing by the kernel does not induce significant nonlinearities. *c* is dependent on the height and width (the integral) of is dependent on the height and width (the integral) of the kernel and was measured here as 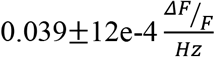. The consistent, negative bias observed in estimating information with 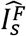 (Figure 2E) would be easy to correct for assuming the *c*factor, and therefore the kernel, were similar across all measured neurons. This point is considered further in the *Guidelines for application of information metrics to functional fluorescence imaging data* section below. We conclude that ground information, as measured by the fluorescence SMGM bits per second metric 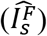, is transformed into different units and is linearly scaled by a factor *c*dependent on the height and width of the kernel.

The amplitude (height) of the change in fluorescence can vary across indicators and conditions. The height of the kernel, given a constant kernel width, should linearly scale *c* and the error in estimating information with 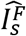. To explicitly test this prediction, we simulated an additional 5,000 fluorescence traces with kernels of varying height (0-3 ΔF/F, Figure 2I-K), but that maintain the same shape and width (from the GCaMP6f kernel), and then measured the percent error in estimating information with 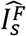. As observed above for the GCaMP6f example (Figure 2 E-G), the percent error in estimating information with 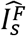 shows little dependence on ground truth information (Figure 2I-K). However, as a function of the height of the kernel, the percent error (averaged over all ground truth information values) in estimating information with 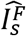 is fit well with an increasing linear function (intercept = −99.8±0.42%, intercept p<<0.01, slope 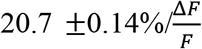, slope p<<0.01, R^2^= 0.80; Figure 2I-)J. Over the wide array of available functional fluorescent indicators in use today (Figure 2H), this leads to differences in error due to differences in transient height of the indicator used alone. For the indicators shown in Figure 2H, there is an average height of 0.603±0.10 SD Δ*F*/*F*: the error spans from −95.8% for the kernel height reported for gCamp6f (0.19 Δ*F*/*F*) to −88.2% for gCamp7f (0.56 Δ*F*/*F*). It should be noted that fluorescence (ΔF/F) is always reported here as a fractional change, not as a percentage (% ΔF/F); if a kernel height of 19 % ΔF/F is used, the units would again change. Thus, as expected, the percent error in estimating information with the SMGM bits per second estimator 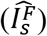 scales linearly with the height of the kernel.

The width of the kernel can vary widely across fluorescent indicators (Figure 2H), with “faster” indicators boasting shorter rise and fall times. The combined effect of a longer rise and fall time is to smooth and delay the AP train; in other words, it acts as a causal low-pass filter. The cutoff period of this low pass filter provides a measurement of the effective width of the kernel [see Methods]. The effect of such differences in kernel shape on the error in estimating information with 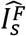 is difficult to measure analytically. We therefore simulated an additional 5,000 fluorescence traces with kernels of different kernel widths (but constant height of the GCaMP6f kernel), resulting in a range of kernel durations (rise times: 1 ms to 1 second, fall times longer than the rise time up to 2 seconds), and then we measured the percent error in estimating information with 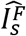. Similarly, as observed above for the GCaMP6f and varying kernel height examples (Figure 2 E-K), the percent error in estimating information with 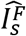 shows little dependence on ground truth information (Figure 2N). Interestingly, the percent error (averaged over all ground truth information values) in estimating information with 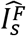 shows a complex nonlinear response as a function of the width of the kernel (Figure 2L-M). The error increases up to a kernel width of ~2.5 seconds, at which point it saturates at ~-85% error. This arises from an interaction between changing the average value of the original AP trace and flattening the average fluorescence map (fi). Over the wide array of available functional fluorescent indicators in use today, this leads to differences in error due to differences in width of the indicator used alone. For example, an average error of −97.1±0.63% were observed for iGluSnfR, the shortest indicator considered here at 0.52 seconds. For gCamp6s, the slowest indicator examined (2.54 seconds), the average was −89.6±4.6%. To estimate the percent errors for these five indicators considering differences in *both* height and duration, we used these 5 kernels to generate mock fluorescence traces from the 10,000 neurons in Figure 2B-G. The resulting distributions, estimated *c* values, and mean and absolute errors can be seen in Figure 2-figure supplement 2. In summary, we conclude that information, as measured by the fluorescence SMGM bits per second metric 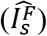, is transformed into different units and is linearly scaled by a factor (*c*) dependent on the height and width of the kernel, with *c* linearly dependent on height and nonlinearly dependent on width. The error induced by these transformations changes substantially over the range of kernel values of the different functional indicators widely used today, and therefore these are important factors to consider when designing and interpreting functional imaging experiments (see *Guidelines for application of information metrics to functional fluorescence imaging data* for further discussion).

### Quantification of the accuracy and precision of the SMGM bits per AP metric using functional fluorescence recordings

The SMGM metric is commonly normalized by the mean rate to obtain a measurement in units of bits per AP. We thus applied the SMGM bits per AP metric 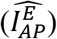 to our mock AP recording traces to verify that they can recover our ground-truth information values. Figure 3A shows three mock neurons with ground truth information values 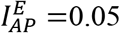, 1.8 and 4.2 bits/AP. When the SMGM bits per AP metric 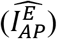 was applied to the AP traces from these example neurons, the information was well recovered, with 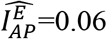, 1.8 and 4.2 bits/AP respectively. The results from these examples also held across the full 10,000 mock neuron library (Figures 3B-D), as a linear fit (y-intercept = 0.087±0.029, intercept p=2.8e-184 bits per second and slope = 0.93 ±0.0010, slope p<<0.01) explained nearly all the variance (R^2^ = 0.99), the average error was −0.071±0.23 bits per AP (3.2±5.9% error) and the absolute error was 0.13±0.21 bits per second (8.1±9% error). However, the data was better fit with a saturating exponential (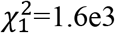, p<<0.01) converging to 5.8 bits/AP as it approached the limit due to the finite bin count. There is a substantial positive bias for the lowest firing rates and smallest number of trials (Figure 2-figure supplement 1E-F) which has been previously well characterized (Treves and Panzeri, 1995). Thus, the SMGM bits per-AP metric 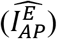 recovers the ground-truth information well using AP recordings (except at the largest ground truth information values), with the primary error coming from finite recording time and variable animal behavior.

**Figure 3.**
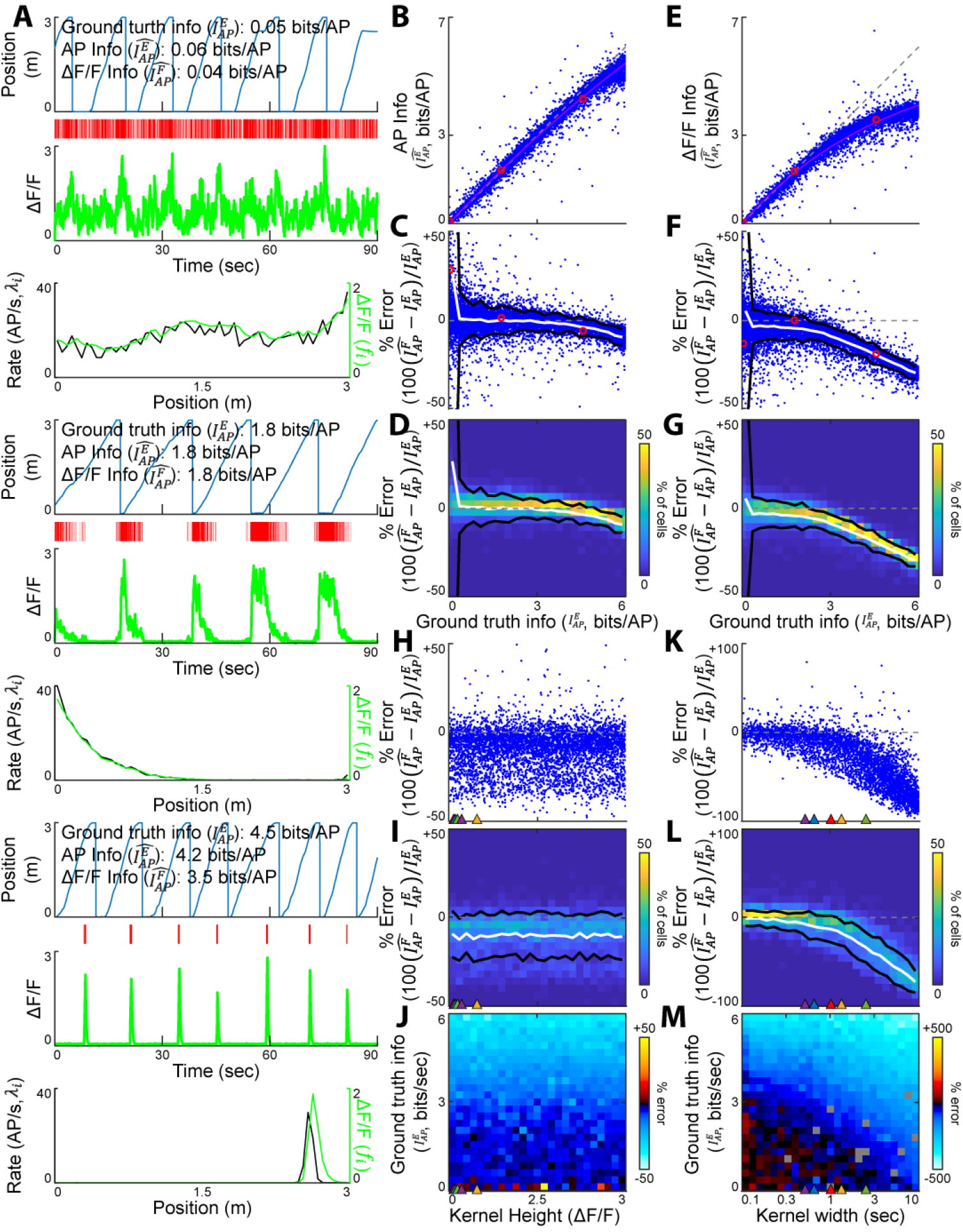
Quantification of the precision of the SMGM bits per AP metric using APs or functional fluorescence recordings. (A) Three representative mock neurons spanning the range of ground truth information values in bits per AP 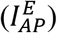. From top to bottom for each: mouse track position vs time, AP raster, fluorescence calcium trace (green), and firing rate map (λ_*i*_, black) and change in fluorescence map (*f_i_*, green). (B-D) The ground truth bits per AP values are well recovered when measured from AP traces. (B) Information measured from AP data using the SMGM bits per AP metric 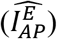 vs ground truth information 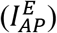. Each dot is a single mock neuron, the gray dashed line is the unity line (perfect measurement). Red circles show the examples in A. (C) Percentage error for the information measurements shown in B. (D) Heat map of percentage error measurements shown in C. Black lines are 2 standard deviations, the white line is the mean. (E-G) Effects of applying the SMGM bits per AP metric to fluorescence traces. (E) Information measured from mock GCaMP6f traces using the SMGM bits per AP metric 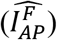 vs ground truth information 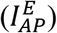. (F) Percentage error for the information measurements shown in E. (G) Heat map of percentage error measurements shown in F. (H-J) The effect of kernel height on estimating ground truth information 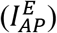 using the SMGM bits per second metric 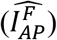. Kernel height for the kernels shown in Figure2H are indicated by colored triangles (H) Percentage error as a function of kernel height (I) Heat map of percentage error measurements shown in H with mean (white) and 2 standard deviations (black). (J) The average percentage error as a function of kernel height and ground truth information in bits per AP 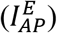. (K-M) The effect of kernel width on estimating ground truth information 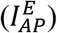 using the SMGM bits per AP metric 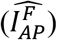. Kernel widths for the kernels shown in Figure 2H are indicated by colored triangles. (L) Percentage error as a function of kernel width (M) Heat map of percentage error measurements shown in L with mean (white) and 2 standard deviations (black). (N) The average percentage error as a function of kernel width.

We next discuss the changes needed to apply the SMGM bits per AP metric 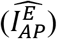 to functional fluorescence traces and explore the implications of these changes. Most simply, the mean firing rate 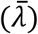 and the mean firing rate in a spatial bin (λ_*i*_) are replaced by the mean change in fluorescence 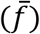 and the mean change in fluorescence in a bin (*f*_i_). Making these substitutions in Equation 5 results in the information as measured by functional fluorescence:

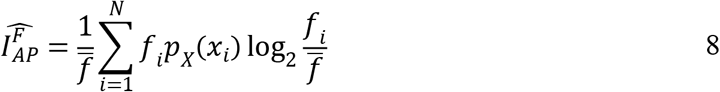

As discussed above, the fluorescence map (*f_i_*) can be approximated as a scaled version of the rate, that is, *f* = *c*λ and 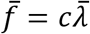. Thus, under this approximation, the *c* factors in Equation 8 cancel, leading to 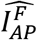 equivalent to 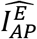, with the same units of bits/AP. This, of course, ignores the fact that the kernel smooths the rate map, leading to a bias in the metric that is difficult to quantify analytically.

We then applied the fluorescence SMGM bits per AP metric 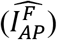 to our 10,000 mock GCaMP6f traces. Figure 3A shows the fluorescence traces generated from three mock neurons with ground truth information of 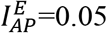, 1.8 and 4.2 bits/AP. When the fluorescence metric 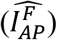 was applied to the fluorescence traces in these examples, the information recovered was 0.04, 1.8 and 3.5 bits/AP, indicating some deviations – especially for the highest information neuron. These results held for the 10,000 mock neuron library (Figure3E-G). At low information values, there was little bias, but at higher information values the information recovered was substantially lower than the ground truth information. The mean resulting error was −0.38±0.58 bits/AP (−9.7±27.8%) and absolute error of 0.39 (12.9±26.4%). This error was better fit with a saturating exponential than a linear fit (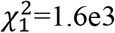, p<<0.01), with the average error less than 5% up to ground truth information of 1.8 bits/AP and less than 10% up to 3.0 bits/AP. At ground truth information values higher than 3 bits/AP, the average error was −1.06±0.595 (−22.5±9.44%) and absolute error was 1.07±0.589 bits/AP (22.6±9.21%). This error persisted even with denser sampling: it remained consistent even at high firing rates and many trials (Figure 2-figure supplement 1E-F). Thus, the indicator induces relatively little error at lower information values (<3 bits/AP), but the smoothing effect of the kernel induces a nonlinear, negative bias to the estimator, particularly at ground truth information values over 3 bits/AP.

Although the height of the kernel can vary between different functional fluorescence indicators (Figure 2H), these height variations linearly scale the fluorescence map. Thus, since 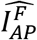 involves normalization by the mean change in fluorescence 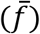, 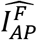 should not depend on kernel height. To explicitly test this prediction, we used the 5,000 fluorescence traces described in the previous section (Quantification of the accuracy and precision of the SMGM bits per second metric using functional fluorescence recordings), with kernels of varying height (0-3 ΔF/F), but that maintain the same shape and width (from the GCaMP6f kernel). Then, we measured the percent error in estimating information with 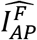 (Figure 3H-J). Unlike for the SMGM bits per second metric, the percent error (averaged over all ground truth information values) in estimating information with 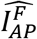 shows little or no dependence on the height of the kernel (p=0.43), but a nonlinear dependence on ground truth information as in Figure 3E-G, with no significant difference in the parameters of the saturating exponential fit (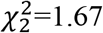, p=0.43). Thus, as expected, the percent error in estimating information with the SMGM bits per AP metric 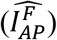 does not vary with the height of the kernel.

With little effect of kernel height on 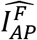, the width of the kernel likely drives biases in the metric. We thus used the 5,000 fluorescence traces generated from a range of different kernel durations (rise times: 1 ms to 1 second, fall times longer than the rise time up to 2 seconds; but constant height of the GCaMP6f kernel) from the previous section (*Quantification of the accuracy and precision of the SMGM bits per second metric using functional fluorescence recordings*), and then we measured the percent error in estimating information with 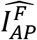 (Figure 3 K-M). Similarly, as observed above for GCaMP6f and the varying kernel height examples (Figure 3E-J), the percent error in estimating information with 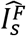 shows a nonlinear dependence on ground truth information (Figure 3E-G). The percent error (averaged over all ground truth information values) showed a nonlinear response as a function of the width of the kernel (Figure 3K-L), with a steep increase in error for kernel widths >~1 second. Even for kernel widths <~1 second, the percent error was strongly dependent on the ground truth information value, with steep increases in error for values >~2.5-3 bits/AP (Figure 3M). Thus, as the kernels gets wider, there is more negative bias at lower and lower information measured. The resulting errors are thus larger for wider kernel indicators: for example, with a kernel width the same as gCaMP6s (2.54s), the error exceeds −17% even at low (<0.25 bits/AP) information, with average errors of −0.86±1.0 bits/AP (−31±19% error) and absolute errors of 0.87±1.0 bits/AP (32.6±16% error). In contrast, with a kernel width the same as iGluSnfR (0.52 seconds), the average error exceeded 5% at 3 bits per AP and 10% at 3.7 bits per AP with a mean error of −0.57±1.00 (−8.0±14%) bits/AP and absolute error of 0.41± 0.56 bits/AP (11±11%). To estimate the percent errors for the five indicators shown in Figure 2H, taking into account differences in *both* height and duration, we used the 5 kernels to generate mock fluorescence traces from the 10k neurons in Figure3B-G. The resulting distributions, mean and absolute errors, and error thresholds can be seen in Figure 3-figure supplement 1.

Since the known information values in our library of 10,000 mock neurons were determined using the SMGM metric, which includes the assumption that neuron firing follows an inhomogeneous Poisson process, we next investigated whether the biases observed between AP and fluorescence metrics (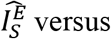 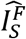 and 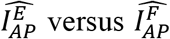; Figures 2 and 3) in our mock neuron datasets were also observed in real neuron recordings (i.e. real spiking that could deviate from Poisson firing).We therefore measured information in a real spiking dataset from hippocampal neurons in rats running on a behavioral track (Chen et al., 2016; Grosmark and Buzsaki, 2016; Grossmark et al., 2016). We generated mock fluorescence traces as we did with simulated AP trains from our mock neurons, compared the information measured from APs versus fluorescence (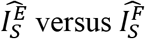 and 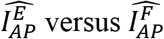) in the real neuron recordings and found that the biases were largely consistent with the simulated mock neuron datasets (Figure 5-figure supplement 1).

In summary, we conclude that ground truth information, as measured by the fluorescence SMGM bits per AP metric 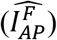, retains the units and insensitivity to height scaling of the electrophysiological metric 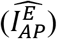, but is nonlinearly biased by the smoothing of the fluorescence map dictated by the width of the kernel. The estimation errors strongly depended on both the width of the kernel and the information value being measured. Since these parameters change substantially over the different functional indicators and different neuron types and behaviors that are commonly used today, they are important factors to consider when designing and interpreting functional imaging experiments (see below for further discussion).

### Nonlinearity introduces further biases

The results presented in the previous two sections rely on the approximation that ΔF/F scales linearly with the firing rate, which is not strictly true in practice (Dana et al., 2018; Éltes et al., 2019; Greenberg et al., 2018). Calcium imaging can be more responsive to bursts of APs rather than isolated spikes, and saturates at high firing rates. As an example for how to examine how nonlinearities between Δ*F*/*F* and firing rate could affect the fluorescence SMGM metrics (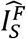 and 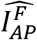), we applied a log-sigmoid nonlinearity (Figure 3-figure supplement 3A) to the 10,000 mock GCaMP6f time-series traces described above, based on the real behavior of GCaMP6f in cultured neurons (Dana et al., 2019, see Methods). While the resulting measurements (Figure 3-figure supplement 3) of ground truth information, as measured by the fluorescence SMGM metrics, are largely consistent with the results observed when using the linear assumption (Figures 2 and 3), some quantitative difference can be seen. Thus, even a relatively simple nonlinearity between ΔF/F and firing rate can add distortions to the amount of information measured using the fluorescence SMGM approach.

### Deconvolution may not be sufficient to eliminate biases

The framework presented here for comparing ground truth information with information measured with the SMGM metrics can be extended to test the efficacy of other strategies for extracting mutual information. In particular, a perfect AP inference method would alleviate the problems associated with applying the SMGM metrics to functional fluorescence recordings. To test the utility of such a strategy in measuring information, we applied a popular deconvolution algorithm, FOOPSI (Vogelstein et al., 2010, see Methods), to the same 10,000 mock GCaMP6f time-series traces described above. Importantly, this deconvolution algorithm (and other available algorithms) does not recover traces of relative spike probability or exact spikes times, but instead produces sparse traces with arbitrary units, that have non-zero values estimating the relative ‘intensity’ of spike production over time (*d*). This signal can be thought of as a scaled estimate of the number of spikes per time bin, and thus the average intensity map will have some similar properties to the florescence intensity maps – that is, we would expect the intensity maps from deconvolution to approximate the relative firing rate scaled by some factor *c*, which has arbitrary units.

We then measured information in these deconvolved*d*-traces using the SMGM metrics (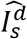 and 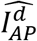), which are identical to (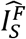 and 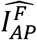), except SMGM is applied to *d*-traces instead of functional fluorescence traces. When using the SMGM bits per second measure (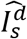, Figure 4A), we found a clear scaling of the ground truth information. The scaling factor was very small (*c*=1.15e-3±1.8e-5 A.U.), resulting in low predicted information (mean error −11.4±6.92 AU, mean % error −99.8±0.38%). This error was significantly larger than when we measured information directly from the florescence traces using 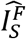 (Figure 2E; 96.0% absolute error, Ranksum p<<0.01; c=0.0390). It is worth noting that the deconvolved trace *d* can be arbitrarily scaled, so in a sense this error is arbitrary. However, these are the results from the scaling chosen by a widely used deconvolution algorithm and the large error emphasize that the scale of *d* can have a large effect on the bits per second measure 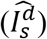.

**Figure 4.**
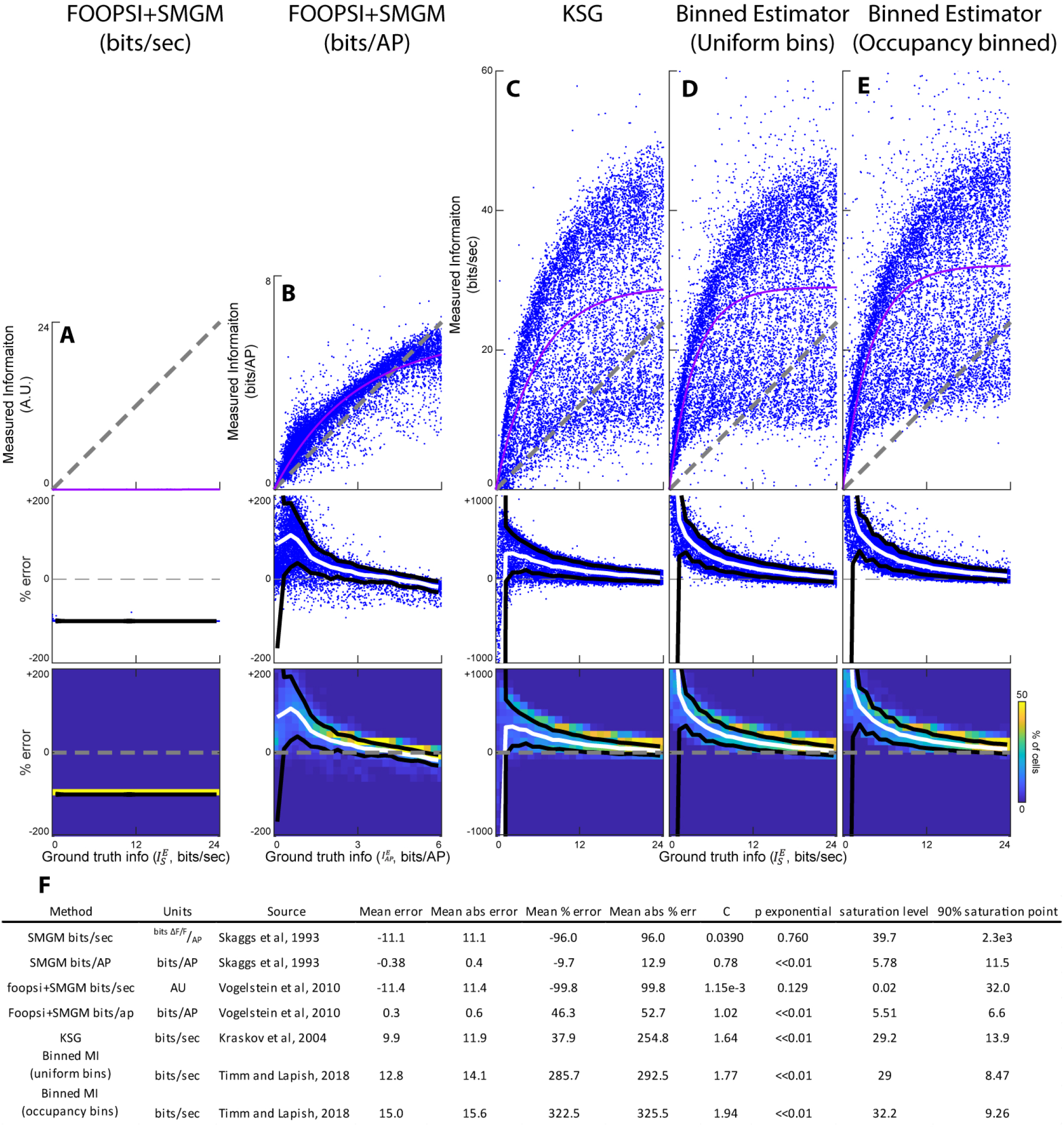
Alternative techniques for measuring mutual information from functional fluorescence traces. (A-E) (Top) Information measured from mock GCaMP6f traces vs ground truth information. The gray line is the unity line, the pink line is the best fit saturating exponential. (Middle) Percentage error for the information measurements shown on top. (Bottom) Heat map of percentage error measurements shown in middle. (A) FOOPSI deconvolved traces using the SMGM bits per second metric 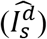. (B) FOOPSI deconvolved traces using the SMGM bits per AP metric 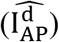. (C) The KSG measure applied to GCaMP6f traces. (D) The Binned Estimator applied to GCaMP6f traces using uniform bins. (E) The Binned Estimator applied to GCaMP6f traces using equal occupancy bins. (F) Table of summary statistics for each measure. P exponential is the p-value from the chi^2^test used to determine if a saturating exponential fit is better than a linear fit for the measured vs ground truth information plots.

Assuming that the intensity map of the deconvolved *d*-traces are a scaled version of the true rate maps, we could measure information using the SMGM bits per AP metric 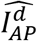 without changing units (Figure 4B). Compared to the SMGM bits per AP metric applied to florescence 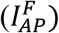, on average there was some reduction in the nonlinearity at higher ground truth information 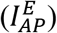 values when using 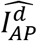, resulting in linear fits closer to the unitary line (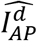 slope = 1.02 ± 0.0027, R^2^ = 0.76 versus 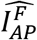 slope = 0.78±0.0011, R^2^=0.93). However, information measured with 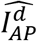 was still better fit with a saturating exponential (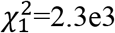, p~0) converging to a saturation value of 5.51 bits per AP (compared to 5.78 for 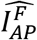), as expected since the algorithm is not expected to resolve spikes at orders of magnitude shorter timescales than the kernel. This resulted in a positive bias at lower levels of ground truth information. For ground truth information values below 3 bits per AP, the average error for 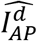 was 0.556±0.50 bits per AP (68.9±104%) compared to −0.060±0.13 bits per AP (2.75±0.36%) for 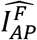. For ground truth information values above 3 bits per AP, the average error for 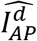 was −0.127±0.76 bits per AP (−1.0±16.8%) as compared to −1.04±0.59 bits per AP (−0.22±9.4%) for 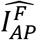. Overall, there was more error when the SMGM bits per AP metric was applied to deconvolved data compared to when applied directly to fluorescence traces (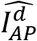 mean absolute error 0.60±0.47 bits/AP (52.7±89.1%) vs 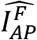 was 0.13±0.21 bits per second (8.1±9% error), Ranksum p<<0.01). Thus, when comparing the recovery of ground truth information from functional fluorescence traces using either direct application of the SMGM metrics (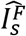 and 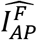) or the application of the SMGM metrics to deconvolved z-traces (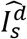 and 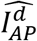), we found better recovery using the direct application approach (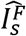 and 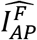).

### The KSG and Binned Estimators are poor estimators of MI in functional florescence data

In addition to SMGM, the KSG and Binned Estimation metrics have been developed for estimating mutual information between variables. These other two metrics produce information measured in bits per second, so they are only comparable to the SMGM bits per second estimator 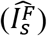. The KSG metric uses the kth nearest neighbor distances between points in the neural response and behavioral variable space to estimate information (Kraskov et al., 2004). The Binned Estimation metric uses discrete bins to estimate the full multidimensional joint probability distribution (*p*_(*X,Y*)_ in Equation 1) to estimate mutual information (Timme and Lapish, 2018). These two metrics estimate information across time samples and therefore are dependent on firing rate like the SMGM bits/sec metric considered above. Further, the Binned Estimator is sensitive to the precise method used for data binning, and thus we have used two commonly applied binning methods: uniform and occupancy based bins.

We applied the KSG, Binned Estimator (Uniform bins), and Binned Estimator (Occupancy binned) (see *Methods*) to the same 10,000 mock GCaMP6f time-series traces and behavioral data used to assess the SMGM approach. These methods all behaved similarly when applied to our simulations (Figure 4C-E), so they will be discussed together here. The information values measured by these techniques correlated with ground truth information in bits per second 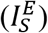. Interestingly, unlike the SMGM bits per second metric (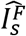, Figure 2), the KSG and Binned Estimator results were better fit with a saturating exponential than with a linear fit (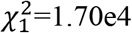, 3.12e4, and 3.14e4 respectively, p~0). The KSG and Binned Estimator methods overestimated the information at lower ground truth 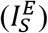 values and saturated quickly at higher values. For ground truth information 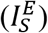 values below 10 bits per second, the mean absolute errors were 10.3±7.82 bits per second (465±4,390%), 22.2±12.6 bits per second (821.7 ±3,353%), and 24.2±14.0 bits per second (911±3,926%) for the KSG, Binned Estimator (Uniform bins), and Binned Estimator (Occupancy binned) respectively (Figure 4F). This is in comparison to the 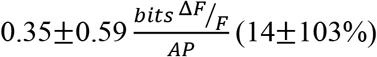 mean absolute error found using the SMGM bits per second metric 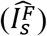 for ground truth information values less than 10 bits per second. For ground truth information 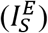 values greater than 10 bits per second, the mean absolute errors were 13.2±8.66 bits per second (85.0±64.4%), 25.4±16.2 bits per second (165.2±118.3%) and 28.9±17.34 bits per second (186.3±124.9%) for the KSG, Binned Estimator (Uniform bins), and Binned Estimator (Occupancy binned) respectively. This is in comparison to the 0.97±1.15 bits per second (5.85±6.73%) mean absolute errors found using the SMGM bits per second metric 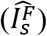 for ground truth information values greater than 10 bits per second. Over the full range of ground truth values, mean absolute errors of 11.9±8.42 bits per second (254.8%), 23.5±14.8 bits per second (456.7%) and 26.7±16.1 bits per second (509.0%) were found for the KSG, Binned Estimator (Uniform bins), and Binned Estimator (Occupancy binned) respectively – an order magnitude larger than the 2.4±2.97 AU (22±27% error) error seen using the SMGM bits per second metric 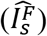. As a control, we applied the Binned Estimators to AP traces and compared the estimated information to the ground truth information to verify that the large errors observed (Figure 4 D,E) were caused when the estimators were applied to fluorescence data (rather than simply a difference between the Binned Estimator, which do not rely on a Poisson firing assumption, and the ground truth information established using SMGM, which does rely on a Poisson firing assumption). We found the errors when applying the Binned Estimators to AP traces were relatively small (Figure 4-figure supplement 2, mean absolute error 2.72±3.38 bits per second (41.1%) and 2.70±3.33 bits per second (41.0%) for the uniform and occupancy based binning respectively). Therefore, when comparing the recovery of ground truth information from functional fluorescence traces using either the SMGM metric 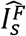 or the KSG and Binned Estimator metrics, we found better recovery using the SMGM approach 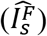.

### Guidelines for application of information metrics to functional fluorescence imaging data

Taken together, the above results suggest that across the information metrics applied directly to functional fluorescence traces, the SMGM metrics provide the most reliable and interpretable information measurements. We thus suggest the following guidelines for use and interpretation of the SMGM metrics as applied to fluorescence mutual information metrics (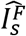 and 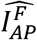) defined in Equations 7 and 8.

The SMGM bits per second metric 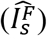 is likely attractive to imaging researchers because the units suggest that precise knowledge of AP numbers and times are not required for its use. However, there are several challenges when applying the SMGM bits per second metric to functional fluorescence imaging data. First, the substitution of the change in fluorescence map (*f*) for the AP firing rate map (λ_*i*_) introduces a change in units, from bits per second to 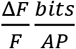, which is difficult to interpret and relate back to bits per second. Second, the transformation of AP firing rate to change in fluorescence can be approximated by a *c* scaling factor (*f*= *c*λ), which is measured in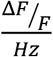, a quantity that is unknown *a priori*. If *c* is not consistent between the neurons of a population of interest, then the information values will be scaled differently and cannot be directly compared (Figure 2). Since *c* is dependent on the width and height of the indicator response to a single AP (the kernel), it can vary from neuron to neuron based on difference in indicator expression level, intracellular calcium buffering, and many other factors (Aponte et al., 2008; Greenberg et al., 2018; Helmchen and Tank, 2015; Park and Dunlap, 1998). More research will be needed to measure these parameters (e.g. Chen et al., 2012, 2013a), and thus *c*, across neurons. Some results suggest that there may be non-trivial amounts of variability within a populations of neurons (Éltes et al., 2019; Greenberg et al., 2018). With the impact of *c* on the SMGM bits per second metric, and the possible variability of *c* across a population of neurons, how can researchers properly extract useful measurements of information using 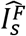 ?

Guideline 1: First, we note that *if* experimental measurements reveal small and acceptable variations in *c* across the neurons of interest, then the information values derived from 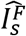 can be normalized by this factor to recover information values in units of SMGM bits per second (independent of Δ*F*/*F*) that can be compared across neurons.

Under the assumption of a consistent kernel, approximations for *c* for common indicators can be found in Figure 2-figure supplement 2.

Guideline 2: Further, given small variations in *c* across the neurons of interest, the ratio of 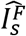 between neurons in the population provides a meaningful metric for comparisons. For example, such ratios could be used to divide a population of neurons accurately into groups based on their information values (e.g. three quantiles of information) or compare the information values between different functional subtypes of neurons (e.g. between place and non-place cells).

Guideline 3: The metric can still be useful even if experimental measurements reveal large and unacceptable variations in *c* across the neurons of interest, or if experimental measurements of *c* do not exist. In such cases, since it is reasonable to assume that *c* is consistent in the same neuron over time, comparisons across the same neuron can provide meaningful insights by using a ratio of 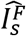 measured (from the same neuron) across different conditions. For example, quantifying the neuron by neuron ratio of 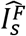 between different behavioral states or conditions of an animal or task, such as between goal-vs non-goal-directed running down a linear track, would allow researchers to make conclusions such as the following: “The population of neurons in region Z carries X+/−Y times more information during goal-directed than non-goal-directed running.”

Therefore, we conclude that with careful consideration of the (known or unknown) variability of the fluorescence response kernel (*c*), 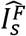 can be used to extract useful measurements of information, either direct measurements of information across a population of neurons (with known and similar *c*), ratios of information between different neurons of a population (with known and similar *c*) or differences across different conditions within the same neuron (with *c* unknown or different across neurons).

The SMGM fluorescence bits per AP metric 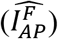 results in the same units as the AP based metric 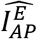, and therefore may provide imaging researchers with information values that are relatively easy to interpret. However, this similarity in units is somewhat misleading since the number and timing of APs are not directly measured with functional fluorescence traces and the asymmetric and relatively slow dynamics of fluorescence indicators leads to shifting and smoothing of the AP rate map (λ_*i*_). This issue can have the effect of inducing a significant negative bias in information measurements, especially at high information values and with functional indicators with wider kernels (Figure 3). This is the most important factor to consider when determining how researchers can properly extract useful measurements of information using 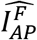. The shifting and smoothing of the AP rate map by fluorescence effectively leads to crosstalk between adjacent spatial bins. Therefore, it is critical to consider the size of the spatial bins in relation to the spatial shift and smoothing induced by the indicator (effectively the kernel plotted in space, rather than time, using the animal’s average running velocity to transform from time to space). It is reasonable then to counteract the spatial shift and smoothing effect by using larger bin sizes, but this only works up to a point since larger bins limit the maximum amount of information possible to measure and may negatively bias the information values near this upper limit, even for AP based recordings (Figure 3B-D). Researchers could potentially optimize the recovery of ground truth information by appropriately selecting bin size for a particular indicator (see Figure 3-figure supplement 2 A-B).

In practice, using gCaMP6f and the rodent spatial behavior and spatial bin sizes (5cm) used here, our analysis suggests that 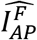 provides reasonable measurements of information for neurons with values up to 3 bits per AP (Figure 3E-G) since this is the point where the absolute error exceeds 10% (comparable to the mean absolute error when measuring information from AP data (8.4%)). Equivalent thresholds for other common indicators are shown in Figure 3-figure supplement 1. The error is exacerbated by slower indicators and thus more accurate measurements of information will result from using the fastest, narrowest kernel indicators available, assuming signal to noise and detection efficiency are comparable across the different width indicators.

Guideline 4: We conclude that with careful consideration of the size of the spatial bins in relation to the spatial shift and smoothing induced by the indicator, 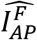 can be used to extract useful measurements of information, most accurately for neurons with < 3 bits/AP under recording conditions similar to those considered here.

Previous research quantifying information in bits per AP using 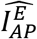 have found that the majority of neurons carry information in this range (< 3 bits/AP) (Bourboulou et al., 2019; Knierim et al., 1995; Lee et al., 2006; Markus et al., 1995; Poucet and Sargolini, 2013)with a few exceptions (Ji and Wilson, 2007). Although these levels are dependent on the number of bins and bin dwell time, 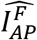 should be widely applicable to quantifying information throughout the brain during behavior.

### Example: application of information metrics to functional fluorescence imaging data from hippocampus during spatial behavior

In this section we demonstrate use of the above guidelines for proper application and interpretation of the SMGM fluorescence mutual information metrics (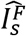 and 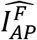) defined in Equations 7 and 8. We applied these metrics to functional fluorescence recordings (gCaMP6f) from pyramidal neurons in CA1 of the hippocampus acquired during mouse spatial behavior.

CA1 neurons expressing gCaMP6f (viral transfection, *Camk2a* promoter) were imaged with two-photon microscopy through a chronic imaging window during mouse navigation along a familiar 1D virtual linear track, as described previously (Figure 5A-B, Dombeck et al., 2010; Radvansky and Dombeck, 2018; Sheffield et al., 2017). 8 fields of view from 4 mice were recorded in 8 total sessions (recording duration 8.8+/−1.3 minutes, number of traversals/session: 29+/−2.5, 3.6+/−0.3 laps/min, 3 m long track). From these 8 sessions, 1,500 neurons were identified from our segmentation algorithm (see Methods), and analysis was restricted to the 964 neurons that displayed at least one calcium transient on at least 1/3 of the traversals during the session. Among these 964 neurons, 304 (31.5%) had significant place fields and were thus identified as place cells (see Methods), while the remaining 660 (68.5%) did not pass a place field test and were thus identified as non-place cells.

**Figure 5.**
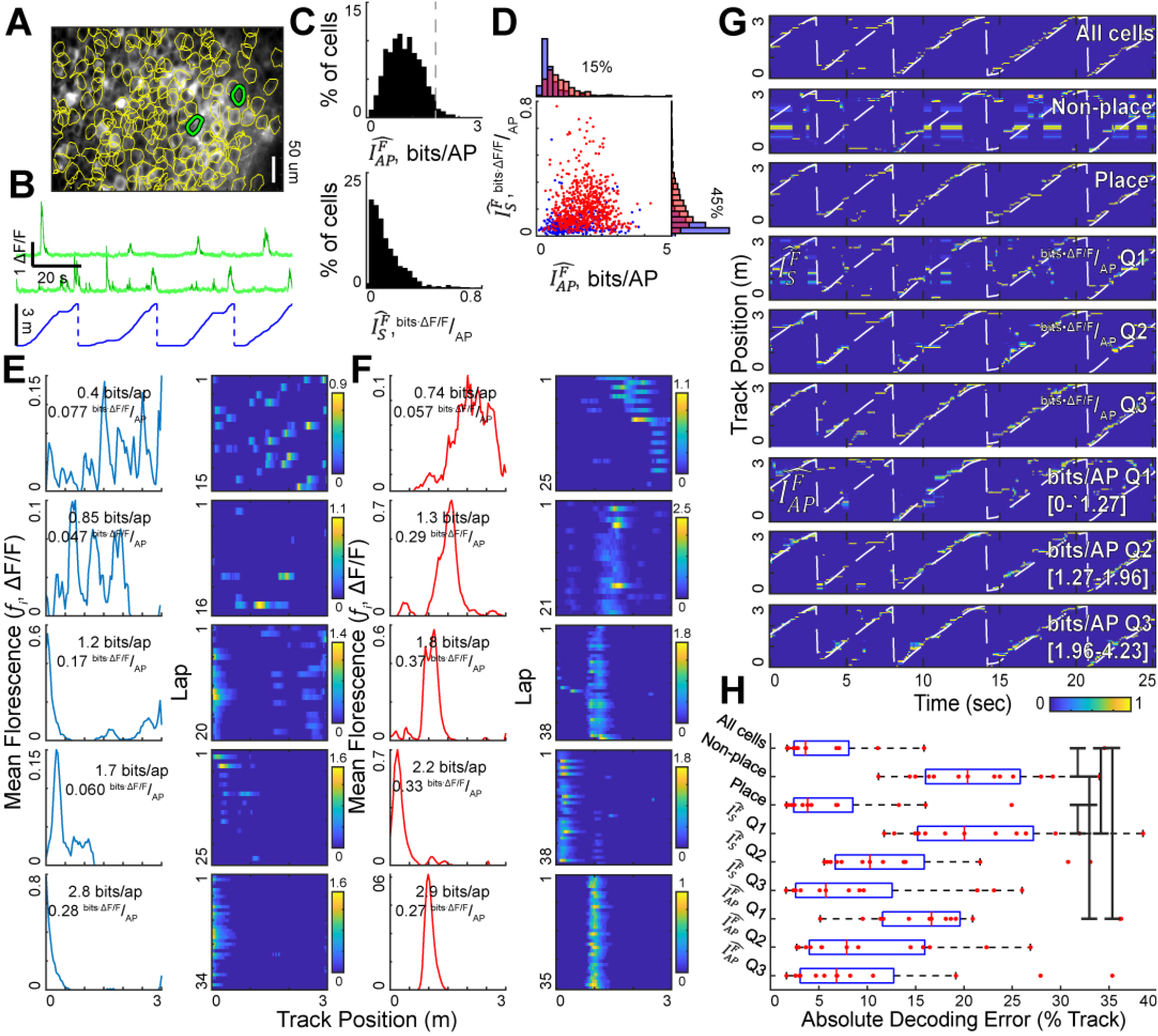
Application of SMGM information metrics to functional fluorescence imaging data from hippocampus during spatial behavior. A.) Example field of hippocampal pyramidal neurons expressing GCaMP6f and imaged during linear track navigation. Active cell ROIs shown in yellow; traces for green cells shown in B. B.) Fluorescence DF/F traces (green) from two neurons in the field shown in A and the track position during the recording (blue). C.). Distribution of information values using the fluorescence SMGM bits per second metric (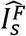,(top) and the fluorescence SMGM bits per AP metric (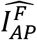, bottom). The gray line indicates the recommended cutoff for reliability using GCaMP6f. D.) Plot of 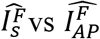 for each neuron. Place cells indicated in red and nonplace cells in blue. E.) Example non-place cells spanning the information ranges shown in C. Spatial fluorescence map (fi) shown on left, and average change in fluorescence per track traversal on right. F.) Same as E, but for place cells. G.) Bayesian decoding of mouse’s track position using different subpopulations of neurons for one example session. From top to bottom: All active neurons, all nonplace cells, and place cells, the first through third quantiles of the SMGM bits per second formulation 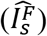, and the first through third quantiles of the SMGM bits per AP formation 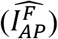. The white dashed line indicates the ground truth position of the animal, the color map indicates the decoded position probability (peak-normalized posterior distribution). H.) Decoding accuracy (absolute position decoding error in units of % of track) pooled over all sessions for each neuron group indicated in G. Black bars indicate significant differences by Holm-Bonferroni corrected rank sum tests (α =0.05).

By applying Equation 7 (using 5 cm sized spatial bins), we found a continuum of spatial information values measured by the fluorescence SMGM bits per second metric 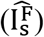 across the 964 CA1 neurons, with an average value of 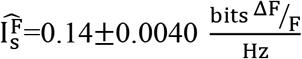 (Figure 5C). The units for 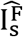 make direct use and interpretation of these values difficult, however, because these recordings were all from pyramidal neurons in a single area, here for illustrative purposes, we will presume that variations in c (discussed above) across the 964 neurons of interest are small and acceptable, with the absolute value of c unknown. This allows for comparisons of information ratios across different subsets of the population. For example, place cells had 2.8±0.20 times more information than non-place cells using the SMGM bits per second metric(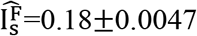 for place vs. 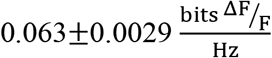 for non-place, Rank sum p=1.7e-63, Figure 5D-F), although there was substantial overlap in information between the populations (see distributions in Figure 5D and individual examples in Figure 5E,F). This also allows for accurate division of the 964 neurons into 3 quantiles based on information values, which we use below for spatial location decoding.

By applying Equation 8 (using 5 cm sized spatial bins), we found a continuum of spatial information values measured by the fluorescence SMGM bits per AP metric 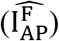 across the 964 CA1 neurons, with an average value of 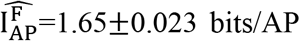 (Figure 5C). The units for 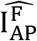 allow for direct use and interpretation of these values, and notably, because most (97%) of the neurons had values less than 3 bits/AP, a mean absolute error of <10% can be assumed across the distribution of SMGM bits per AP values. When applied to the place and non-place cell populations, we found that place cells had higher information than nonplace cells using the fluorescence SMGM bits per AP metric (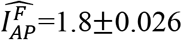 and 1.35±0.042 bits/AP, Rank sum p=4.6e-21, Figure 5D-F). This is consistent with mock fluorescence traces generated from real neuron AP datasets (Figure 5-figure supplement 1B).

As a demonstration of the usefulness of using information metrics to analyze large functional fluorescence population recordings, we explored the accuracy of decoding the animal’s track position using different subsets of neurons. We divided the 964 neurons into 9 groups: All neurons, place cells, non-place cells, three quantiles based on the fluorescence SMGM bits per second metric, and three quantiles based on the fluorescence SMGM bits per AP metric. We then used a Bayesian decoder of the animals’ position (see Methods) separately for each of the 9 neuron groups in each of the 8 sessions (Figure 5G,H). An individual session decoding example can be seen in Figure 5G. We quantified decoding accuracy using the absolute position decoding error (% of track), and pooled this measure across sessions for each neuron group (Figure 5H). The means and standard errors for each group are: All neurons (7.33 +/−2.5 %), place cells (6.97 +/−1.9 %), non-place cells (20.9 +/−1.8 %), SMGM bits per second Q1 (21.9 +/−1.5 %), SMGM bits per second Q2 (13.2 +/−2.4 %), SMGM bits per second Q3 (8.97 +/−2.4 %), SMGM bits per AP Q1 (17.6 +/−2.7%), SMGM bits per AP Q2 (17.8 +/−3.1 %), SMGM bits per AP Q3 (10.4 +/−3.0 %). Interestingly, even the lowest quantile information groups still could be used to determine animal track location to within ~1/5 of the track. This supports the idea that the hippocampal code for space is carried by a large population of active neurons (Meshulam et al., 2016), and not just by a select subpopulation with the highest information or most well-defined tuning curves. As could be expected, place cells encoded the position of the animal better than nonplace cells and better than the lowest quantile information groups (Holm-Bonferroni corrected Rank sum, α=0.05) and neurons in the higher quantiles provided more accurate decoding. Thus, the fluorescence information metrics provide a means to compare the relative contribution of hippocampal neurons with different information values to decoding animal position.

## Discussion

Here, we performed an in-depth simulation study to examine the application of the SMGM bits per second and SMGM bits per AP metrics of mutual information to functional fluorescence recordings. Since these metrics were designed for AP recordings and since functional fluorescence recordings violate some of the assumptions that these metrics are based on, it was unclear if and how the metrics could be used for functional fluorescence recordings. We created a library of ten thousand mock neurons whose AP output carried ground-truth amounts of information about the animal’s spatial location, and by using real behavioral recording data from mice navigating in virtual linear tracks, we simulated the spatial firing patterns of the mock neurons. We then simulated fluorescent calcium responses for each neuron in each session by convolving the AP trains with calcium kernels for different indicators, primarily GCamp6f (though see Supplemental Figures 2 and 3 for results from other indicators), and then added noise.

We then derived fluorescence versions of the SMGM bits per second 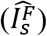 and SMGM bits per AP metrics 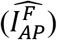 (Equations 7 and 8) and applied them to the fluorescence traces in order to quantify the performance of the metrics for estimating information. We found that ground truth information, as measured by the fluorescence SMGM bits per second metric 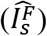, was transformed into different units and was linearly scaled by a factor (*c*) dependent on the height and width of the kernel, with *c* linearly dependent on height and nonlinearly dependent on width. The error induced by these transformations changed substantially over the range of kernel values of the different functional indicators widely used today, and therefore are important factors to consider when designing and interpreting functional imaging experiments. We then found that ground truth information, as measured by the fluorescence SMGM bits per AP metric 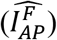, retains the units and insensitivity to height scaling of the electrophysiological metric 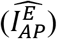, but is nonlinearly biased by the smoothing of the fluorescence map dictated by the width of the kernel. The estimation errors strongly depended on both the width of the kernel and the information value being measured. Importantly, since these parameters change substantially over the different functional indicators and different neuron types and behaviors that are commonly used today, they are important factors to consider when designing and interpreting functional imaging experiments. For example, even for the same indicator, the shape of the kernel is a function of intracellular calcium buffering, indicator concentration, the amount of calcium influx, the efflux rates, background fluorescence and resting calcium concentration, which can all vary across different cells. Additionally, the results presented here rely on the approximation that ΔF/F scales linearly with the firing rate, which is not strictly true in practice. We show in Figure 3-figure supplement 3 that even a relatively simple nonlinearity between ΔF/F and firing rate can add distortions to the amount of information measured using the fluorescence SMGM approach. This relationship between ΔF/F and firing rate can vary across different indicators and, since the Toolbox can be used to vary this relationship, users can further explore this source of bias.

Some of the general features of the relationship between ground truth information and fluorescence SMGM metrics can also be seen using an analytical approximation. For example, if we approximate the rate map as a Gaussian firing field with mean rate 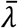, simplify the kernel approximation to a single exponential with falloff τ, assume constant, normalized movement speed *v*, and assume that the convolution between the kernel and the mean rate map is nearly Gaussian, we can approximate the relationship between the bits per second ground truth information 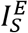 and measured fluorescence information 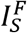 as 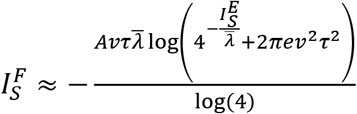 and between the bits per AP ground truth information 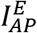 and measured fluorescence information 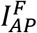 as 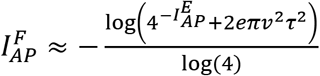. Similar to the numerical simulations, these equations predict that the fluorescence bits per second metric is dominated by a prefactor (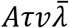 in the analytical case), and that the fluorescence bits per action potential metric saturates at larger information values. Using 1000 values for 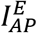 and λ from the same distribution as our simulations, we can approximate a *c* value for 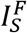 of 0.011 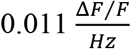 (data not shown). Of course, our numerical solutions provide more accurate approximations for the magnitude of these effects (and for the magnitude of information values themselves) given that they include the more accurate double exponential kernel (leading to a much larger kernel integral compared to a single exponential), signal noise, and the realistic nonstationary speed, position and fluorescence signals.

In our approach, the known information values in our library of 10,000 mock neurons were determined using the SMGM metric, which includes the assumption that neuron firing follows an inhomogenous Poisson process. It is important to remember that the SMGM metric, which has been applied to spiking data extensively over the past few decades, *requires* the use of a Poisson estimate of spiking probability—i.e. the Poisson assumption is built into the original metric. In practice, even spiking data violates this and other assumptions of the SMGM metric since real neurons do not strictly follow Poisson statistics (for example they can display neural hysteresis) and animal behavior is non-stationary. Here we are building from this existing framework and adding and testing whether it is possible to apply the metric to functional fluorescence datasets. Even still, the Poisson assumption could have contributed to some of the biases found when evaluating the fluorescence SMGM metrics with respect to ground truth information. We explored this potential source of bias further using two different analyses. First, in Figure 4-figure supplement 2, we applied the Binned Estimators (which do not rely on a Poisson firing assumption) to AP traces and compared the estimated information to the ground truth information (which was established using the SMGM metric that does rely on a Poisson firing assumption). We found the errors to be relatively small, particularly in comparison to the errors induced by the Binned Estimators when applied to fluorescence traces (Figure 4 D,E). Second, in Figure 5-figure supplement 1, we used a real spiking dataset from hippocampal neurons in mice running on a behavioral track (i.e. real spiking neurons that can deviate from Poisson firing) and generated mock fluorescence traces from the AP traces. When we compared the information measured from the AP traces to the fluorescence traces, we found biases that were largely consistent with those observed in Figure 2 and 3 from our simulated mock neuron datasets. Taken together, these analyses indicate that any biases resulting from the Poisson assumption in the simulation procedure appear to be small, particularly with respect to the biases introduced when AP traces are transformed into functional fluorescence traces. Finally, in the Toolbox, we also include code to generate mock neurons using a binned distribution, avoiding the Poisson assumption of SMGM. Thus users can further explore sources of bias using a different ground truth dataset.

Using our mock fluorescence traces, we also asked if an AP estimation method could relieve the biases in the SMGM metrics. Applying the SMGM bits per second metric 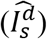 to AP estimation traces from a deconvolution algorithm (FOOPSI) resulted in a low *c* value for recovered versus ground truth information. When the SMGM bits per AP measure was applied 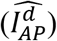, the resulting measurements of information were still nonlinear (compared to 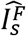), with a positive bias at lower values of ground truth information. Overall, applying FOOPSI to fluorescence traces led to a poorer recovery of ground truth information using SMGM compared to direct application of SMGM to the florescence traces 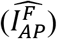. Importantly, this result from deconvolution is only specific to GCaMP6f, and conclusions should not be drawn about other indicators or situations; users will be able to use the Toolbox to explore this area further. We also tested other metrics to measure mutual information directly from the fluorescence time traces (KSG, Binned Estimator (uniform bins), and Binned Estimator (occupancy binned)) and found these alternatives produced highly variable, saturating measurements of recovered versus ground truth information. This was in contrast to the SMGM bits per second measure 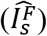 which produced a linearly scaled bias with lower error.

Taken together, we find that the SMGM bits per AP metric can well recover the mutual information between spiking and behavior. The SMGM bits per second metric is scaled such that comparisons should be limited to within populations of well characterized neurons or for within neuron comparisons, e.g. ratios of information across conditions. In general, researchers should use caution when applying measures developed for AP data in fluorescence recordings: there’s no guarantee that the assumptions that support the measures hold for fluorescence data, and this can lead to difficult to interpret and biased results.

## Materials and Methods

### Toolbox and Data Availability

We provide a user toolbox (freely available at https://github.com/DombeckLab/infoTheory), which consists of Matlab functions to generate libraries of model neurons with known amounts of information, to generate spiking or fluorescence time-series from those model neurons, and to estimate neuron information from real or model spiking or fluorescence time-series datasets using the three metrics considered here (SMGM, Binned Estimator, KSG). This toolbox also contains tools to generate mock neurons using a binned distribution, avoiding the Poisson assumption of SMGM. Behavioral data used to generate the random traces is freely available at https://doi.org/10.7910/DVN/SCQYKR.

### Construction of AP trains with known ground truth information

To construct mock neurons with ground truth information, we adapted the differential form of the AP information, in bits per AP (Equation 6). To create a rate map, we first selected an average firing rate and target ground truth information. The mean rate 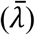 was always between 0.1 and 30 Hz, the information in bits per AP 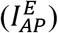 between 0 and 6 bits per AP, and the information in bits per second 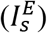 between 0 and 24. To more evenly sample each of these, we first randomly selected the bits per second 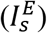 or bits per AP 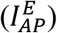 to target. If the information target was in bits per AP, both the information 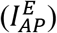 and mean firing rate 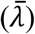 were chosen uniformly. Because the information in bits per second 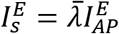, the bits per second information was not uniformly sampled in this case. If the target was to be in bits per second, both the bits per AP 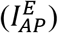 and SMGM bits per second 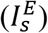 measures were first chosen uniformly. Because the rate 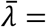 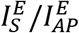, this was not chosen uniformly. This procedure was repeated to maintain the bounds on 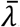, resulting in a non-uniform sampling of information. The final distribution (Figure 1C) was spread acceptably for further analysis.

The rate maps were constructed by spline interpolating across 5 control points with two anchored at each end of the track, and taking the exponential for each point, and then normalizing by the numerically calculated integral (Figure 1A, D). To create a map matching the target information, we began with a random spline. The ‘y’ (relative rate) initial position of each node was chosen from a standard normal distribution and the initial ‘x’ (track position) of the 3 center nodes was chosen uniformly. The nodes were then systematically moved using the MATLAB built in optimizer ‘fmincon’ with constraints preventing the crossing of the center nodes and keeping them on the track, and the ‘OptimalityTolerance’ option set to 0 (Figure 1A). This was accomplished using the ‘genExpSpline’ function' in the toolbox.

We then randomly selected behavioral traces (see Methods *Behavior* section) and concatenated sessions until a total time randomly chosen between 3 and 60 minutes was reached (Figure 1E, F; Figure 2A, Figure 3A). This was accomplished using the ‘loadBehaviorT’ function in the toolbox. The track positions were normalized and used to build a conditional intensity function (CIF) from the rate function above. The CIF was normalized to match an expected mean rate over the entire session, and the MATLAB built-in ‘poissrnd’ function was used to generate AP times, sampled at 1 kHz. The was accomplished using the ‘genSpikeTrain’ function in the toolbox. Finally, the AP times were binned according to the counts within mock imaging frames sampled at 30 Hz.

### Simulated 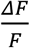 traces

To construct the 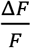 traces (Figure1E,J,K; Figure2A; Figure3A), we first created a single AP response kernel from the peak-normalized sum of two exponentials:

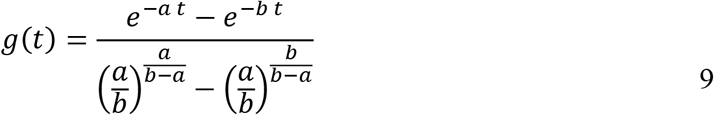

Where *t* is the time since the AP and *a* and *b* are chosen to minimize (1 − *g*(τ_*rise*_))^2^ + (0.5 − *g*(τ_*rise*_ + τ_*fall*_))^2^ where τ_*rise*_ is the rise time in seconds and τ_*fall*_ is the half-fall time in seconds.

Deviations in τ_*rise*_ and τ_*fall*_ from baseline were also measured. The kernel g(t) was then multiplied by the indicator height. The kernel parameters were generated using the ‘fluorescenceKernel’ function, and evaluated using the ‘doubleExp’ function in the toolbox.

The GCaMP6f, GCaMP6s, and jRGECO1a heights, rise and fall times were measured as responses to single APs *in vivo* (Chen et al., 2013; Dana et al., 2019): other kernels (Figure 2H, Supplemental Figures 2 and 3) were approximated from other experiments presented in the references (seen in Table 1).

**Table 1.** Properties of indicator kernels used.

| Kernel | Height<br>$\Delta F/F$ | Rise<br>(s) | Fall<br>(s) | Source |
| --- | --- | --- | --- | --- |
| *gCaMP6f* | 0.190 | 0.042 | 0.142 | (Chen et al., 2013) |
| *jRGECO1a* | 0.164 | 0.041 | 0.207 | (Kalko et al., 2011) |
| *gCaMP7f* | 0.560 | 0.063 | 0.276 | (Dana et al., 2019) |
| *gCaMP6s* | 0.230 | 0.179 | 0.550 | (Chen et al., 2013) |
| *iGluSnfR-A184S* | 0.300 | 0.022 | 0.106 | (Marvin et al., 2018) |

To define the width of the kernel (Figure2L-N, Figure3K-M), we considered the kernel as a low pass filtered version of the APs. If we normalize the filter to mean 1, it has the Fourier transform 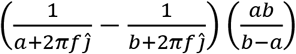. The kernel width was defined as the −3 dB (50%) cutoff period of this filter: 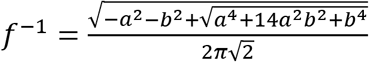. For the simulations with different width kernels, a kernel width was chosen between 0.01 and 10 seconds, a rise time between 0.001 and 1 second, and a fall time between the rise time and 2 seconds. Then, a and b were chosen to minimize the squared error between these three targets using the built in MATLAB optimizer ‘fminsearch’.

White noise with a standard deviation of 0.15 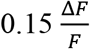 was then added to the mock fluorescence traces.

### Nonlinearity

In our linear simulations used throughout this work, the fluorescence kernels associated with a fast sequence of action potentials were approximated to sum linearly. In real cultured neurons, a summation nonlinearity has been observed such that sequences of action potentials do not generate a linear summation in ΔF/F (Dana et al., 2019). To simulate this nonlinearity, the 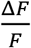 trace was then further transformed as:

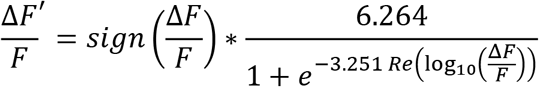

This equation was arrived at by fitting the measured responses in Dana et al, 2019, Figure 2C, which can be compared to the nonlinearity used here (Figure 3-figure supplement 3A).

### Deconvolution

Deconvolution was performed using the previously described FOOPSI algorithm (Friedrich et al., 2017; Vogelstein et al., 2010). The regularization coefficient was set at 0.02154, which maximized the correlation between the deconvolved trace and the true spike train in a random sample of 500 simulated traces: all other parameters were optimized for each trace. Because the example regularization coefficient provided by Friedrich et al, 2017 was 2.4, we also measured information values at 100 different values for the regularization coefficient between 0 and 3; this had little effect on the measured information (Figure 4-figure supplement 1).

### KSG estimator

The previously described second KSG estimator (Kraskov et al., 2004) was used using the 5^th^ nearest neighbor distance.

### Binned estimators

The binned mutual information estimators were used (Timme and Lapish, 2018). The activity trace was divided into 10 bins, either evenly across the span of the activity (uniform binned) or variably so the bins contained the same number of samples (occupancy binned). Position was similarly divided into 60 bins.

### Bayesian decoding

The Bayesian decoder used here (Figure 5 G,H) was adapted from a previously described method (Zhang et al., 1998). Decoding was performed on the likelihood that a significant transient occurred in a time frame, trained on the first 80% of the session and tested on the last 20%. The session was divided into Δ*t*=0.1 second bins. The conditional likelihood that an animal is in position *x_i_* given the number of active frames during a time window (*n*) is 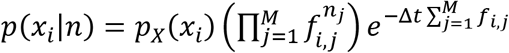, where *p*_x_(*x_i_*) is the (marginal) probability that the animal is in the *i*^*th*^ spatial bin during a time sample, *f_i_,_j_* is the average rate of significant frames by the *j*^*th*^ neuron in the *i*^*th*^ spatial bin, *n_j_* is the number of significant frames observed during the time window in neuron *j*, and *M* is the total number of neurons. The decoded position was selected as the one with maximum conditional likelihood.

### Animals

10 to 12 week old male C57BL/6 mice (20-30g) were individually housed under a reverse 12 hr. light / dark cycle, all experiments were conducted during the dark phase. All experiments were approved by the Northwestern University Animal Care and Use committee.

### Behavior

We used a previously described virtual reality set-up and task (Heys et al., 2014; Sheffield et al., 2017; Sheffield and Dombeck, 2015); some of the behavior sessions used here has previously appeared in these studies. Briefly, water scheduled, head fixed mice were trained to run on a cylindrical treadmill down a 3m virtual track to receive a water (4 ul) reward at the end of the track, and were subsequently teleported to the beginning of the track after a 1.5 s delay. Behavioral sessions were included if the animal ran at least 20 laps containing a continuous 40 cm run for which the velocity was over 7 cm/sec during a 5-30 minute session.

### Mouse surgery and virus injected

We performed population calcium imaging of CA1 neurons as described previously(Sheffield et al., 2017; Sheffield and Dombeck, 2015). Briefly, 30 nL of AAV1-SynFCaMP6f (University of Pennsylvania Vector Core, 1.5*10^13 GC/ml) was injected through a small craniotomy over the right hippocampus (1.8 mm lateral, 2.3 mm caudal of Bregma; 1.25 mm below the surface of the brain) under Isoflurane (1-2%) anesthesia. 7 days later, a hippocampal window and head plate was implanted as described previously (Dombeck et al., 2010).

### Two-photon imaging

Imaging was performed as previously described(Sheffield et al., 2017; Sheffield and Dombeck, 2015). Scanimage 4 was used for microscope control and acquisition (Pologruto et al., 2003). Time series movies (1024 or 512×256 pixels) were acquired at 50 Hz. A Digidata1440A (Molecular Devices) with Clampex 10.3 synchronized position on the linear track, reward timing, and the timing of image frames.

### Image processing, ROI selection and calcium transient analysis

Images were processed as previously described (Sheffield et al., 2017; Sheffield and Dombeck, 2015), with minor modifications. Briefly, rigid motion correction was performed using cross correlation as in (Dombeck et al., 2010; Miri et al., 2011; Sheffield and Dombeck, 2015), but here using a Fast-Fourier transform approximation on the full video. ROIs were defined as previously described (Mukamel et al., 2009) (mu=0.6, 150 principal/independent components, s.d. threshold = 2.5, s.d. smoothing width=1, area limits = 100-1200 pixels). 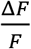 traces were generated by normalizing around the 8^th^percentile of a 3 second sliding window. Significant transients from both experimental and mock fluorescence traces were selected by comparing the ratio of amplitudes and durations of positive to negative going transients with a false positive rate <0.01% (Dombeck et al., 2010). Mock traces used the histograms generated from the mock gCaMP6f traces (Figure 2-3) or from the specific matching indicator traces (Supplemental Figures 2,3): experimental data histograms were built separately. All subsequent analyses were run using these significant transients.

### Behavior analysis

The mean virtual track velocity was defined as the total virtual track distance covered during the session divided by the total duration of the session; slow and stop periods were included in this metric. All other analyses were restricted to long running periods, where the animal exceeded a virtual track velocity of 4 cm/s and ran continuously for at least 40 cm.

### Defining place fields

Place fields were defined by first creating the spatial fluorescence intensity map (*f_i_*) with the 300 cm track divided into 60, 5 cm bins. This map was smoothed via a 3 bin boxcar. Transients identified during run periods were shuffled in order and to random intervals to create 1000 bootstrapped intensity maps. Candidate fields were defined as regions of the original fluorescence map with values greater than 99% of the bootstrapped maps. Fields were then retained if they were between 20 and 120 cm wide: significant place cells retained at least one field that satisfied these criteria.

---

**Figure 2-figure supplement 1.**
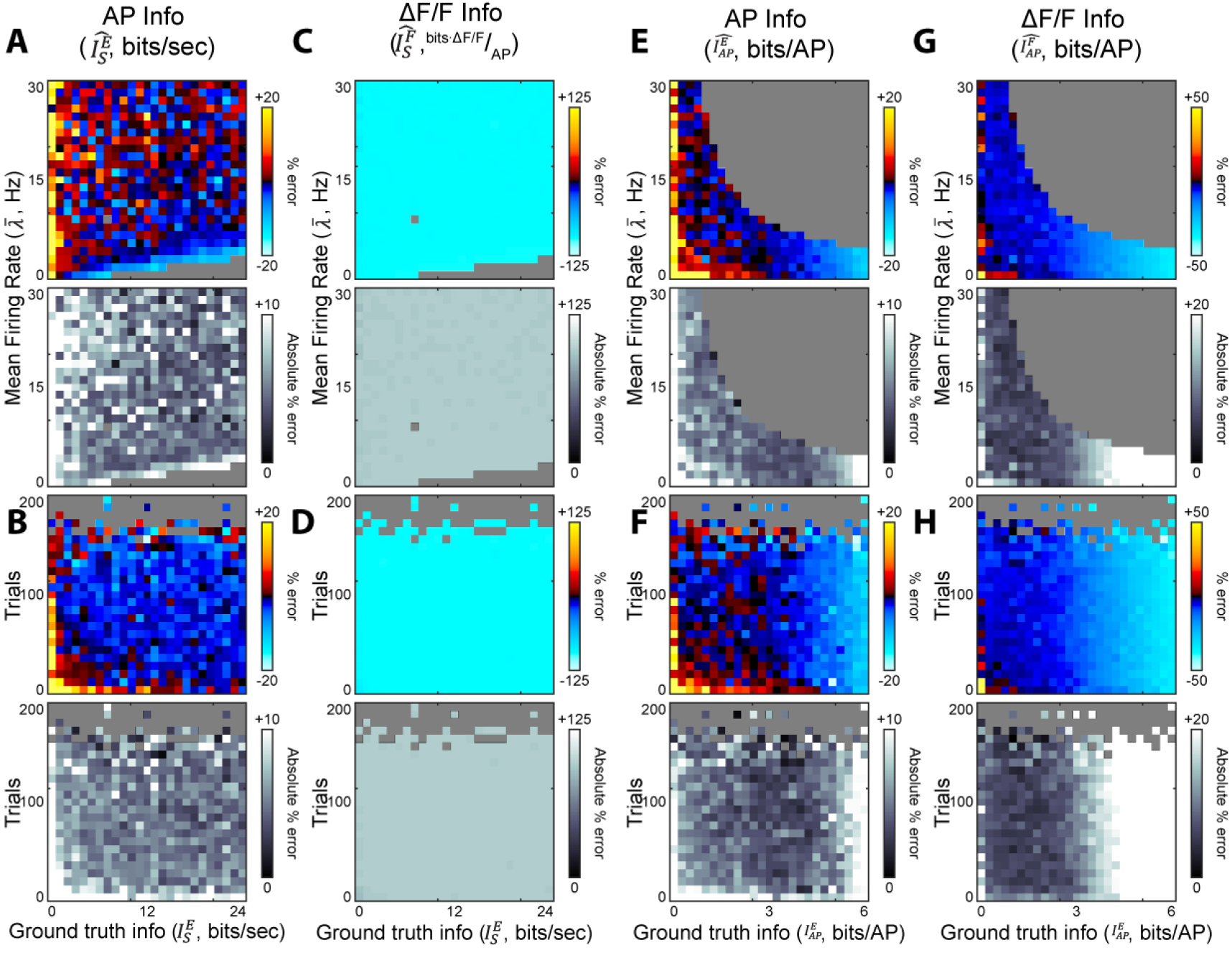
Effect of recording density on information metrics. A. The mean error (top) and absolute error (bottom) between the ground truth information 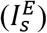 and measured information in SMGM bits per second from mock APs 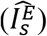 as a function of the mean firing rate 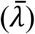. B. As A, but as a function of number of laps. C-D. As A-B, but for measurements from mock fluorescence traces. E-F. As A-B, but for the SMGM bits per AP metric. G-H.) As C-D, but for the SMGM bits per AP metric.

**Figure 2-figure supplement 2.**
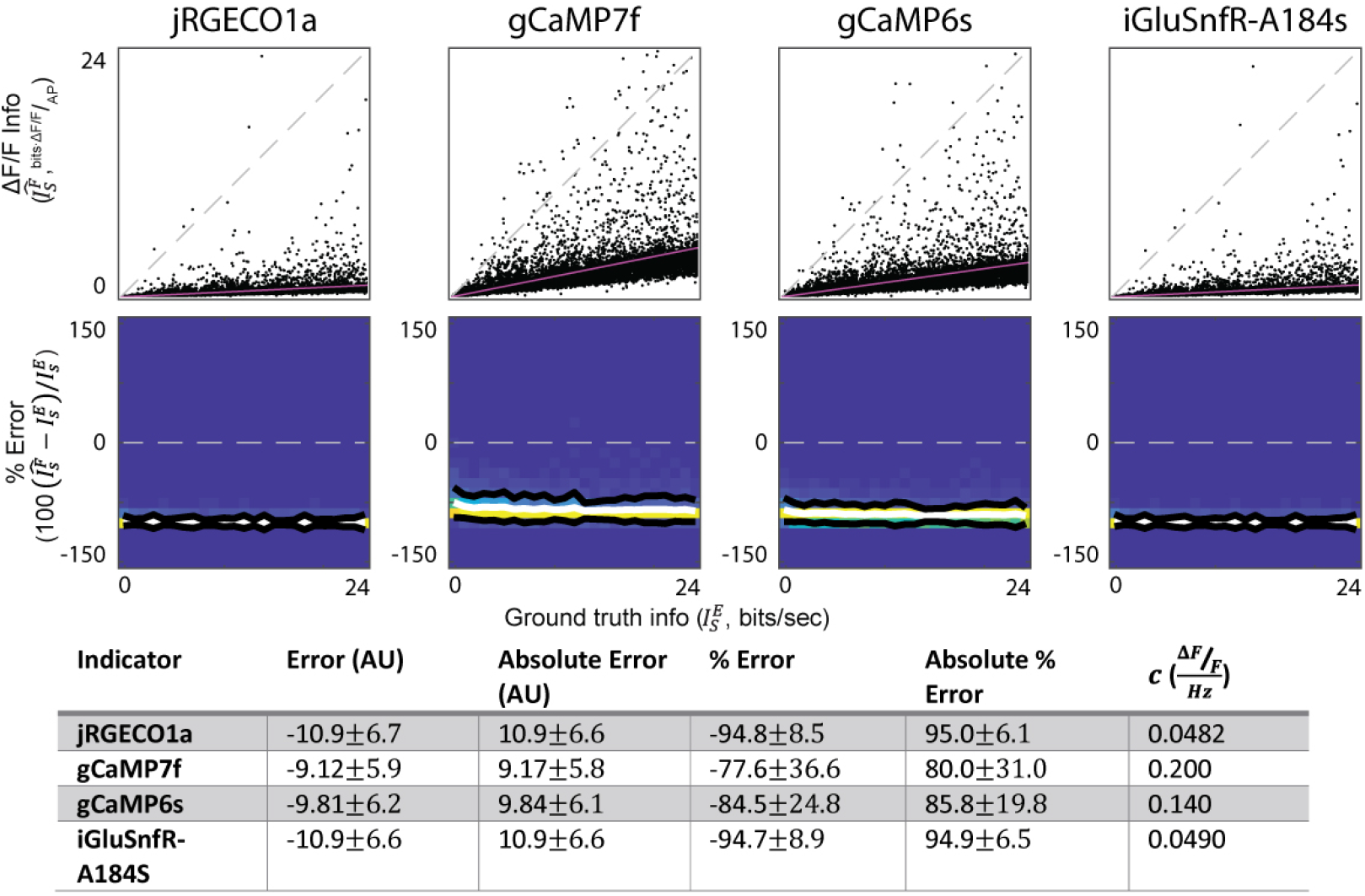
Effects of applying the SMGM bits per second metric to fluorescence traces from different common functional indicators. Top: Information measured from mock traces using the SMGM bits per second metric 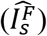 vs ground truth information 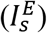. Each dot is a single mock neuron, the gray dashed line is the unity line (perfect measurement), the pink line is the line of best fit. Middle: Percentage error density plots. The white line is the mean, the black are ± one standard deviation. Bottom: Summary statistics and estimates for the scaling factor *c*.

**Figure 3-figure supplement 1.**
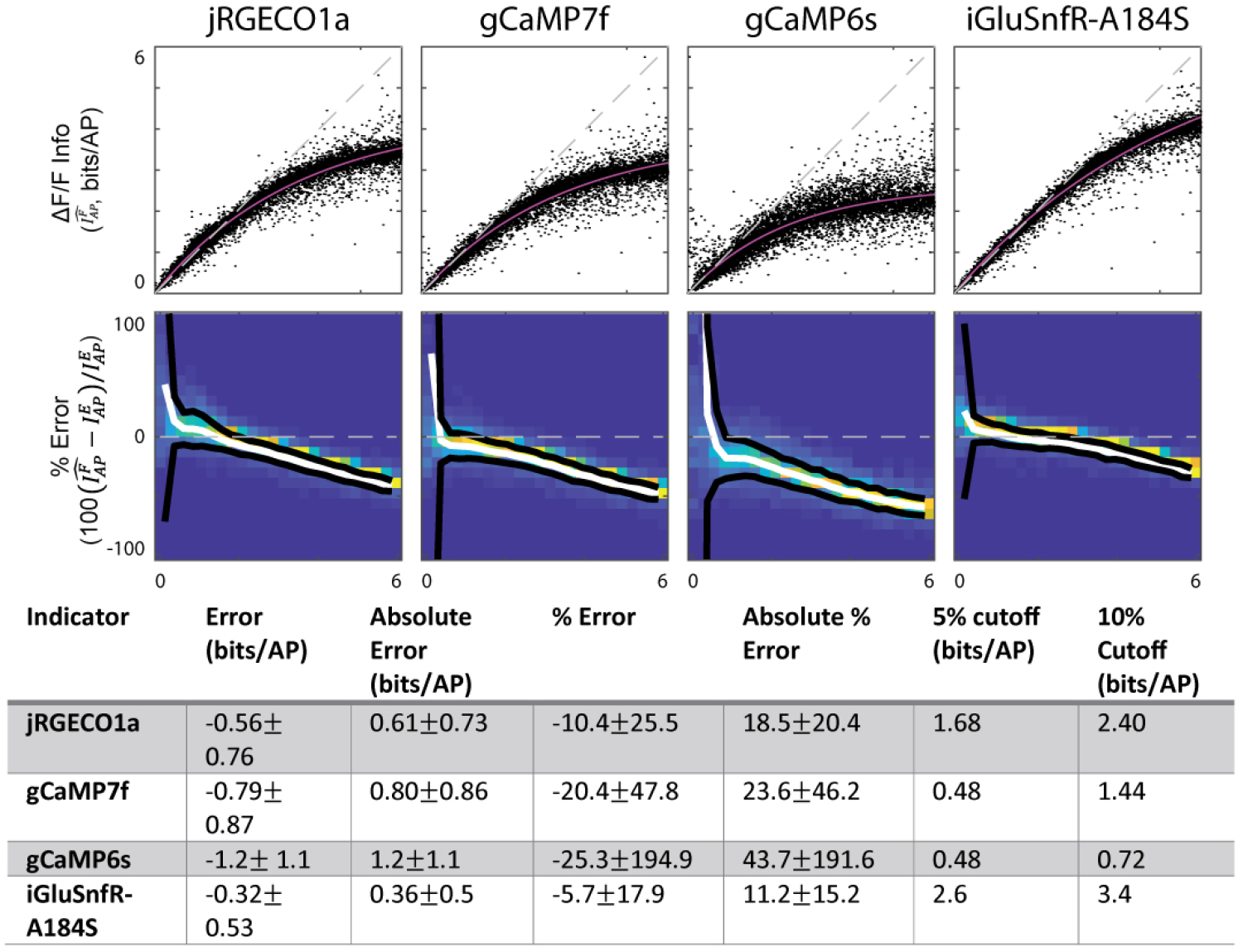
Effects of applying the SMGM bits per AP metric to fluorescence traces from different common functional indicators. Top: Information measured from mock traces using the SMGM bits per AP metric 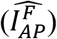 vs ground truth information 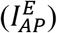. Each dot is a single mock neuron, the gray dashed line is the unity line (perfect measurement), the pink line is the exponential fit. Middle: Percentage error density plots. The white line is the mean, the black are ± one standard deviation. Bottom: Summary statistics and percent error cutoffs.

**Figure 3-figure supplement 2.**
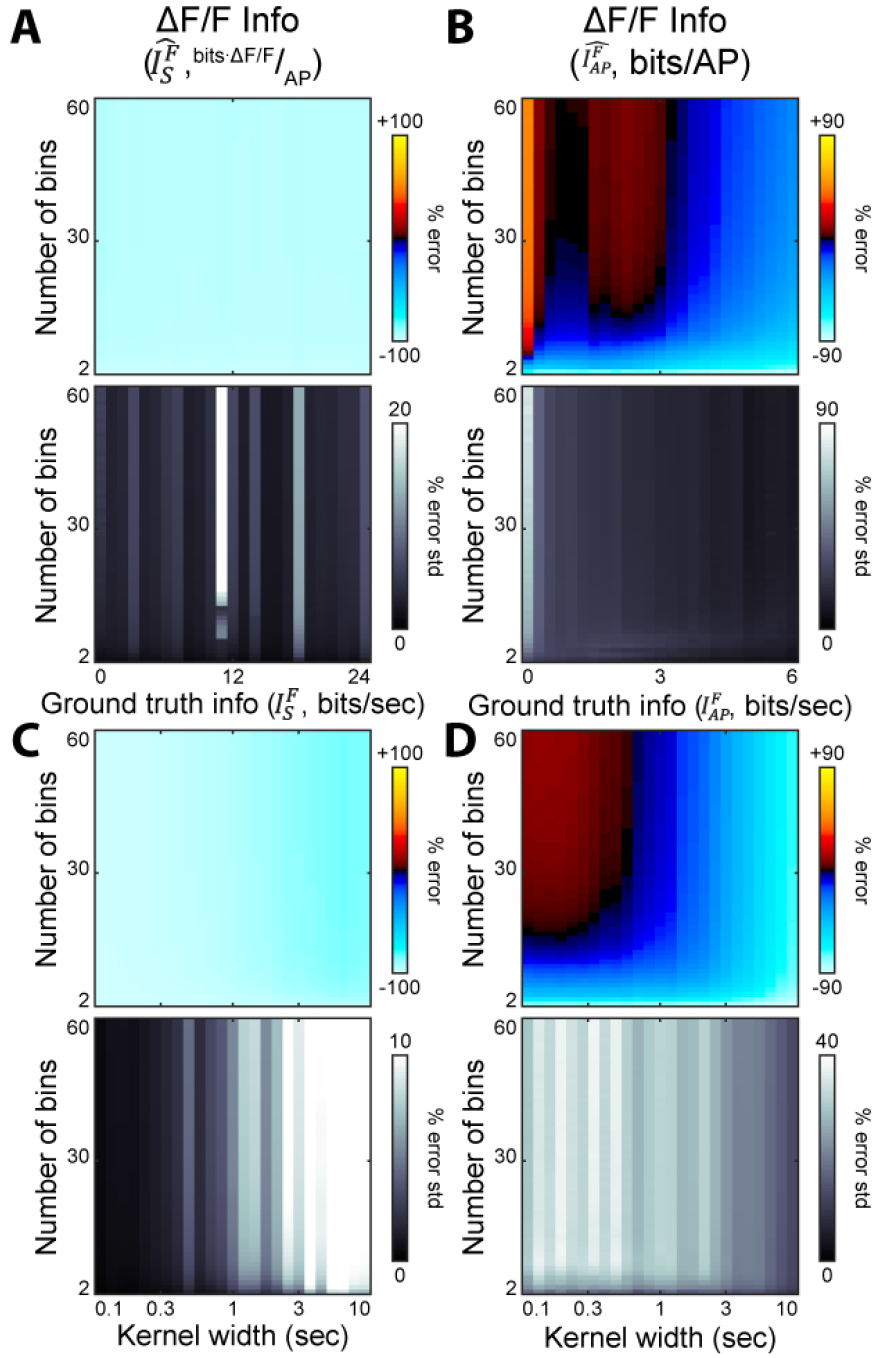
Effect of number of bins on the SMGM metrics. A. The mean percentage error (top) and the standard deviation (bottom) for the bits per second measure 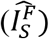 applied to 10,000 mock gCamp6f traces, with ground truth information on the X-axis and 2-60 bins (3m track) on the Y-axis. B. As A, for the bits per AP measure 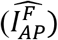. C. The mean percentage error (top) and the standard deviation (bottom) for the bits per second measure 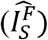 applied to 20,000 mock fluorescence traces with differing kernel width. Kernel width is on the X-axis, number of bins is on the Y-axis. D. As C, but for the bits per AP measure 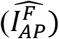.

**Figure 3-figure supplement 3.**
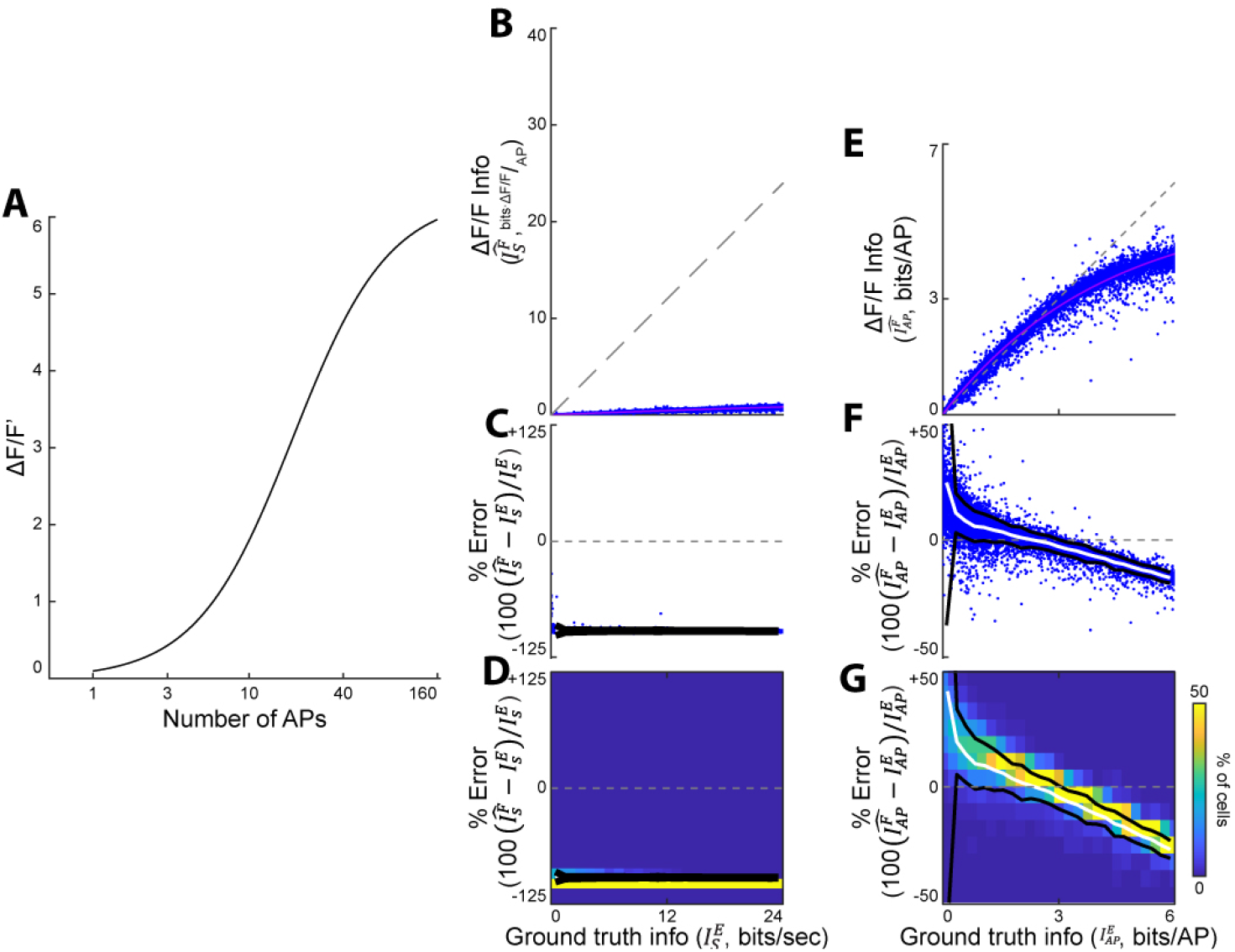
Effects of a sigmoid nonlinearity between ΔF/F and firing rate. Here we applied a log-sigmoid nonlinearity to the 10,000 mock GCaMP6f time-series traces and then measured information using the fluorescence SMGM metrics. (A) Nonlinearity applied to AP-to-florescence trace transformation (B) Information measured from AP data using the SMGM bits per second 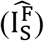 vs ground truth information 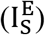. Each dot is a single mock neuron, the gray dashed line is the unity line (perfect measurement). (C) Percentage error for the information measurements shown in B. (D) Heat map of percentage error measurements shown in C. Black lines are 2 standard deviations, the white line is the mean. (E-G) As B-D, but for the bits per AP measured 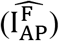 versus the ground truth information in bits per AP 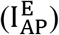.

**Figure 4-figure supplement 1.**
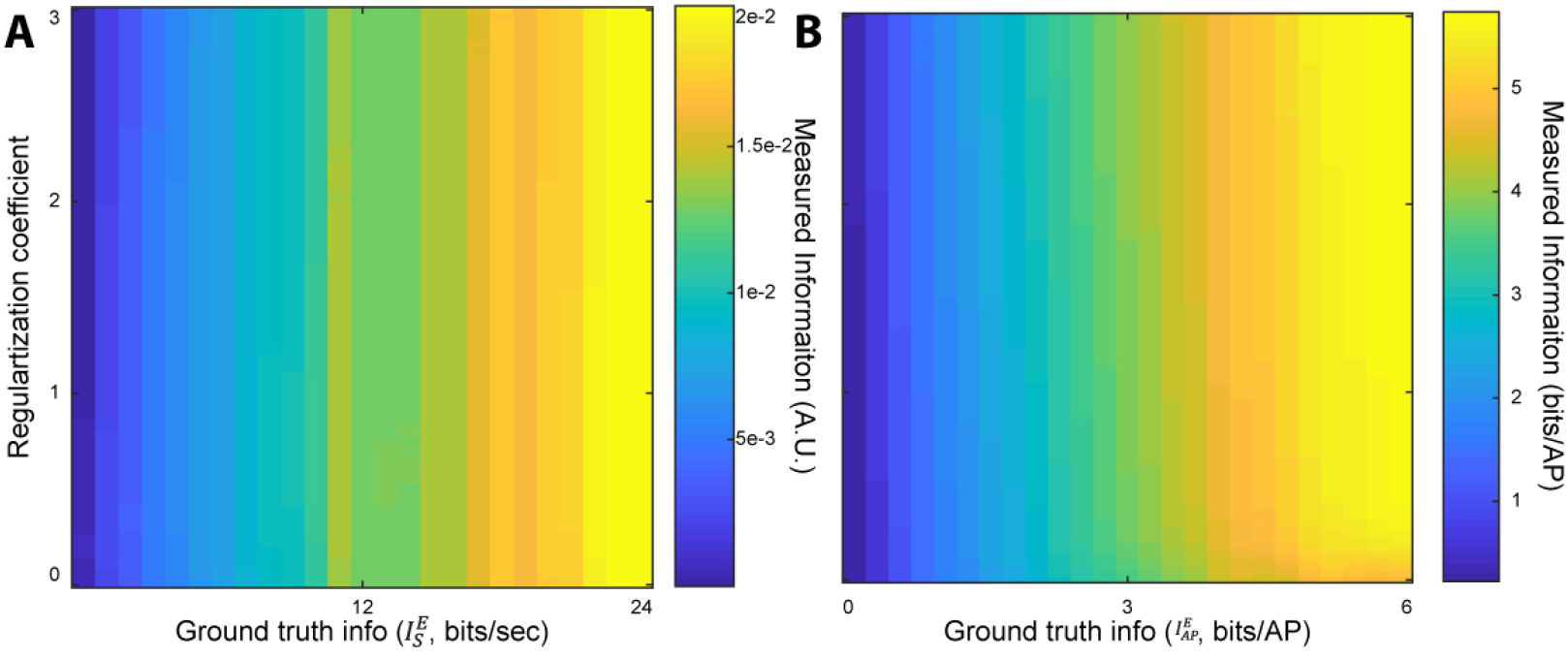
Effect of changing the regularization coefficient in deconvolution (Friedrich et al., 2017; Vogelstein et al., 2010) on the measured information. (A) The measured information using the fluorescence bits per second metric applied to 10,000 mock gCamP6f traces using regularization coefficients between 0 and 3. (B) As A, but for the fluorescence bits per AP metric.

**Figure 4-figure supplement 2.**
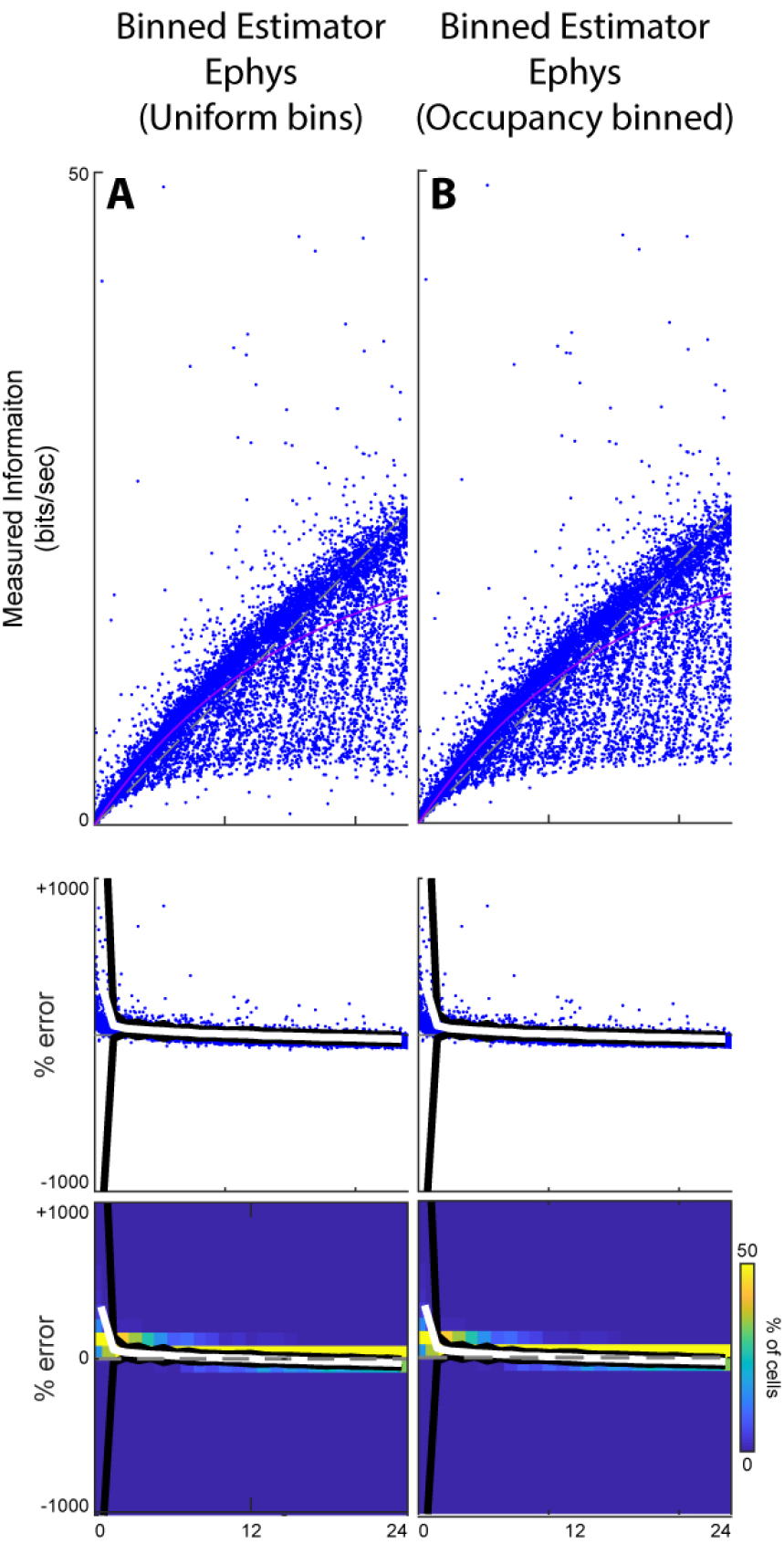
The binned estimator applied to AP traces and then compared to ground truth information. (Top) Information measured from mock AP traces vs ground truth information. The gray line is the unity line, the pink line is the best fit saturating exponential. (Middle) Percentage error for the information measurements shown on top (same scale as shown in Figure 4). (Bottom) Heat map of percentage error measurements shown in middle. (A) The Binned Estimator applied to AP traces using uniform bins. (B) The Binned Estimator applied to AP traces using equal occupancy bins.

**Figure 5-figure supplement 1.**
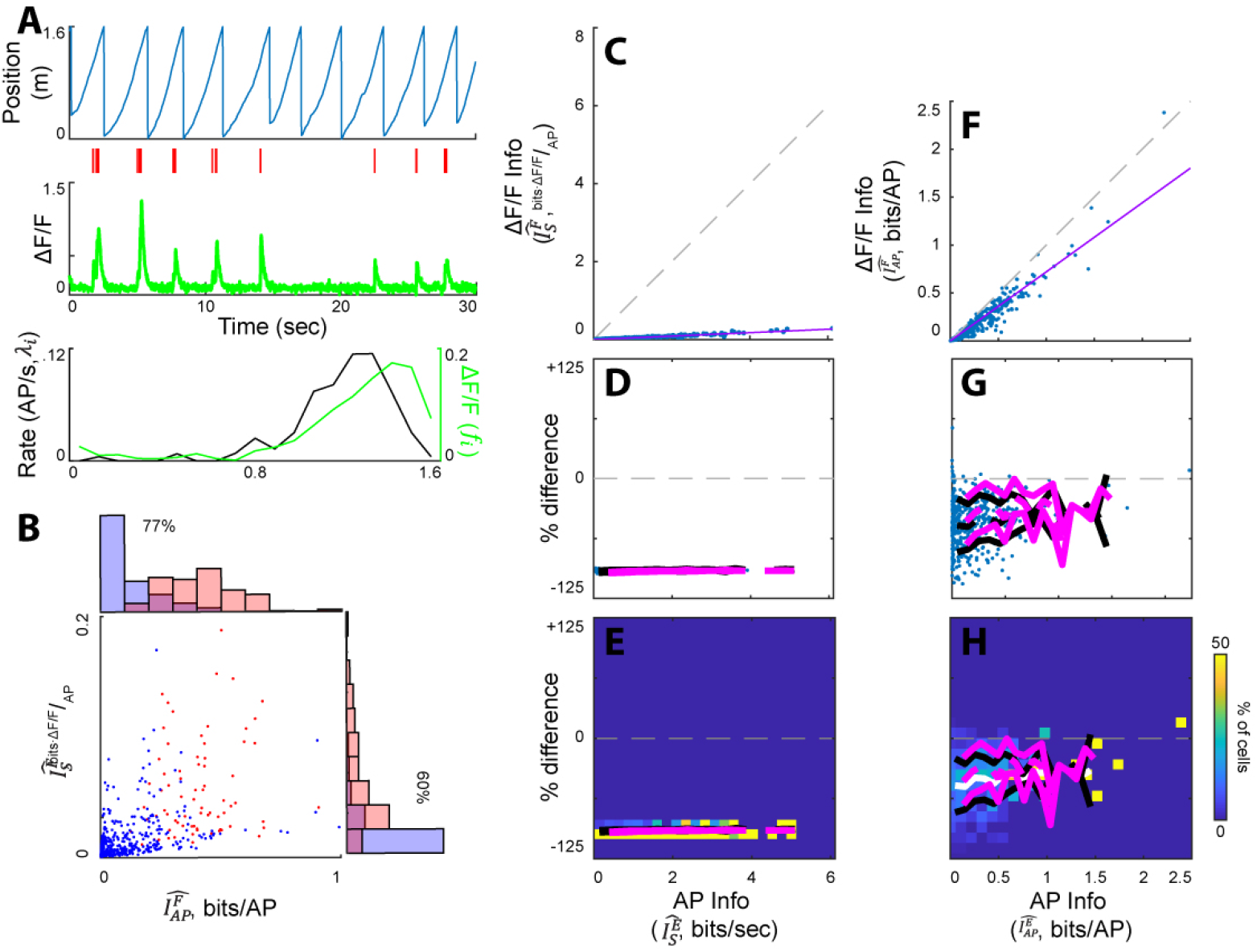
The SMGM estimators as applied to real AP data from a real spiking dataset from hippocampal neurons in mice running on a behavioral track (Chen et al., 2016; Grosmark and Buzsaki, 2016; Grossmark et al., 2016). (A) Example real place cell. From top to bottom: rat track position vs time, real AP raster, mock fluorescence calcium trace generated from real AP trace by convolving APs with GCaMP6f kernel and adding noise (green), and firing rate map(*λ*_*i*_, black) and change in mock fluorescence map (*f_i_*, green). (B)Plot of 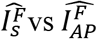 for each neuron. Place cells indicated in red and nonplace cells in blue. (C) The SMGM bits per second metric applied to the real AP traces 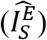 versus the mock fluorescence traces 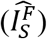 (generated from the real AP traces). (D) The percentage difference between the SMGM bits per second metric applied to the real AP traces 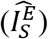 and the mock fluorescence traces 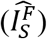. The mean and standard deviation are indicated in black. In magenta, the mean and standard deviation of 5,000 of the mock neuron traces seen in Figures 2 and 3, sampled to have the same firing rates as the real neurons and shortened to the same mean session duration. (E) Density plot for the data shown in D. (F-H) As C-E, but for the bits per AP metric (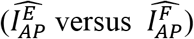).

